# Interpreting graph neural networks reveals microbial association patterns in inflammatory bowel diseases

**DOI:** 10.64898/2026.10.27.762128

**Authors:** André F. Cunha, Edir Vidal, Maria Barranco-Altirriba, Txell Amigó, Juliana G. Assis, Helder I. Nakaya, Alberto Santos

## Abstract

Traditional machine learning models trained on gut microbiome abundance profiles can distinguish individuals with inflammatory bowel disease (IBD) from non-IBD individuals. However, their interpretation typically considers taxa in isolation, making it difficult to understand how associations between taxa contribute to predictions. Here, we inferred individual-specific microbial association networks from longitudinal metagenomics data of 106 individuals in HMP2/iHMP and used them to train and interpret graph neural network (GNN) models, which we compared with abundance-based models such as Random Forest (RF). Although RF achieved higher predictive performance (AUPRC 0.59 vs 0.446), interpreting the GNNs revealed that their predictions relied on a densely connected module of co-abundant Bacteroidales species. Unlike the taxa prioritized by RF, these species were not differentially abundant between conditions. In an independent cohort, this module carried most of the predictive signal, and removing its associations reduced performance to near chance. These results show that coordinated association structure, rather than abundance shifts alone, is a feature of the IBD gut microbiome, and that interpreting GNNs can uncover such patterns that complement abundance-based analyses.

## Introduction

A microbiome is defined as a complex community of microorganisms — the microbiota — along with their functional genetic material, occupying a distinct ecological niche such as the human gut (Berg et al., 2020). The human gut microbiome has been extensively studied in recent years due to its role in human health and disease. This is driven largely by its immediate proximity to the intestinal barrier, which modulates the transport of metabolites into the bloodstream (Van Hul et al., 2024). Specifically, the microorganisms from the gut microbiota are responsible for the degradation of complex molecules including polysaccharides derived from the host’s diet, the synthesis of bioactive compounds such as short-chain fatty acids and vitamins, and competition for resources with pathobionts (i.e., normally harmless resident microbes that can promote inflammation under altered conditions (Chow et al., 2011)) that could otherwise proliferate and disrupt gut homeostasis (Van Hul et al., 2024). This dynamic symbiotic host-microbiota relationship is one of the main reasons why humans and other eukaryotes are often considered “meta-organisms”, which means that they cannot be decoupled from their own microbiomes (Simon et al., 2019; Bosch and McFall-Ngai, 2011).

Changes in the structure and composition of the human gut microbiota have been associated with conditions of different domains, such as gastrointestinal diseases, metabolic conditions, and neurological disorders (Gomaa, 2020). Within the gastrointestinal domain, inflammatory bowel diseases (IBD) serve as a primary example of microbiota imbalance. Specifically, IBD comprises chronic conditions characterized by relapsing and remitting inflammation in the gastrointestinal tract (Ramos and Papadakis, 2019). It manifests mainly as two clinical forms with distinct inflammation patterns: Crohn’s disease (CD), which can affect any area from mouth to anus with uneven, transmural inflammation, and ulcerative colitis (UC), which is restricted to the colon and rectum with superficial mucosal inflammation (Ramos and Papadakis, 2019). The pathogenesis of IBD is not yet fully resolved, but numerous studies indicate that these diseases are caused by an interplay of genetic, environmental, and immunological mechanisms that compromise gut homeostasis (Caruso et al., 2020; Ramos and Papadakis, 2019). One of the main features of IBD is the disruption of the homeostatic host-microbiota relationship, leading to dysbiosis, an alteration of the microbiota composition associated with immune dysregulation and mucosal inflammation (Caruso et al., 2020; Honda and Littman, 2012). Evidence suggests that individuals with IBD have higher abundance of aero-tolerant bacterial clades associated with inflammatory activity and lower abundance of anti-inflammatory clades when compared to non-IBD individuals (Lloyd-Price et al., 2019). Common findings also indicate both a reduction in overall taxonomic diversity and a decrease in the abundance of species belonging to the Firmicutes phylum (Nishida et al., 2018). However, it remains unclear whether dysbiosis contributes to the onset of IBD or arises as a consequence of the disease (Caruso et al., 2020; Sartor and Wu, 2017).

The emergence of metagenomics as a culture-independent environmental sequencing method has enabled the investigation of microbiome links to various diseases, including IBD (Mardanov et al., 2025). Driven by the rapid growth of high-throughput sequencing technologies, its popularity has led to a vast expansion of publicly available metagenomic datasets archived in global repositories, such as the NCBI Sequence Read Archive (SRA) (Leinonen et al., 2010) and EMBL-EBI Mgnify (Richardson et al., 2023), as well as curated catalogues like curatedMetagenomicData (Pasolli et al., 2017) and Metalog (Kuhn et al., 2026), ultimately facilitating large-scale analytical and modeling efforts.

Computational studies analyzing metagenomic data have provided much of the existing insight into gut microbiome patterns in IBD, particularly through the search for microbial markers (Stafford et al., 2022). To unravel the alterations in microbial composition during IBD dysbiosis, these studies frequently utilize machine learning combined with model interpretation methods to identify key taxa that separate IBD from non-IBD individuals (Lee et al., 2025, 2023; Bakir-Gungor et al., 2022; Liñares-Blanco et al., 2022). However, gut microbiome dynamics are shaped not only by shifts in abundance distributions but also by underlying ecological relationships within the microbial community (Liu et al., 2022; Venturelli et al., 2018; Faust and Raes, 2012). For instance, emerging evidence in IBD suggests that interactions between specific species may be linked to alterations in host transcriptional profiles and mucosal immunity (Rice et al., 2022). Consequently, evaluating isolated taxa independently risks oversimplifying the complex ecosystem reconfigurations that might define IBD.

Methods designed to infer microbial association networks (MANs) — such as SPIEC-EASI (Kurtz et al., 2015) and SparCC (Friedman and Alm, 2012) — provide a new analytical dimension for metagenomic studies. Microbial associations represent statistical dependencies between taxa, typically inferred from abundance profiles across samples where species consistently co-fluctuate or exhibit inverse trends (Matchado et al., 2021). Rather than assuming independent behavior, these tools analyze non-random co-occurrence patterns to reconstruct networks that approximate underlying ecological relationships. Inferring MANs enables the identification of systemic properties of the microbiome that would not be detectable solely through the isolated observation of taxonomic abundances (Faust and Raes, 2012). Therefore, interpreting models built on microbial associations rather than independent variables can reveal systemic markers that offer dual value: mechanistic model explainability alongside biologically contextualized insights into the condition.

In this study, we inferred individual-level MANs from longitudinal gut metagenomes and used them to train and interpret graph neural networks (GNNs) for distinguishing IBD from non-IBD individuals. Traditional models trained on abundance tables achieved higher predictive performance, but interpreting the GNNs revealed network-level features that Random Forests did not prioritize. The predictions relied on a densely connected module of co-abundant Bacteroidales species that were not differentially abundant between conditions, and ablation experiments showed that this module carried most of the predictive signal in an independent cohort.

## Results

### Overview of the discovery and validation datasets

We used the publicly available HMP2/iHMP metagenomic dataset (Lloyd-Price et al., 2019) to generate individual-level MANs, train, and interpret the resulting GNNs. This dataset (hereafter referred to as the discovery dataset) consists of 1,627 longitudinal samples from 130 individuals (classified as non-IBD, CD, or UC), with a varying number of samples per individual across the cohort (Fig. 1A). To minimize the inference of spurious individual-level MANs, we excluded data from individuals with fewer than seven longitudinal samples (Fig. 1A, dashed line), which reduced the analysis to 106 individuals.

**Figure 1:**
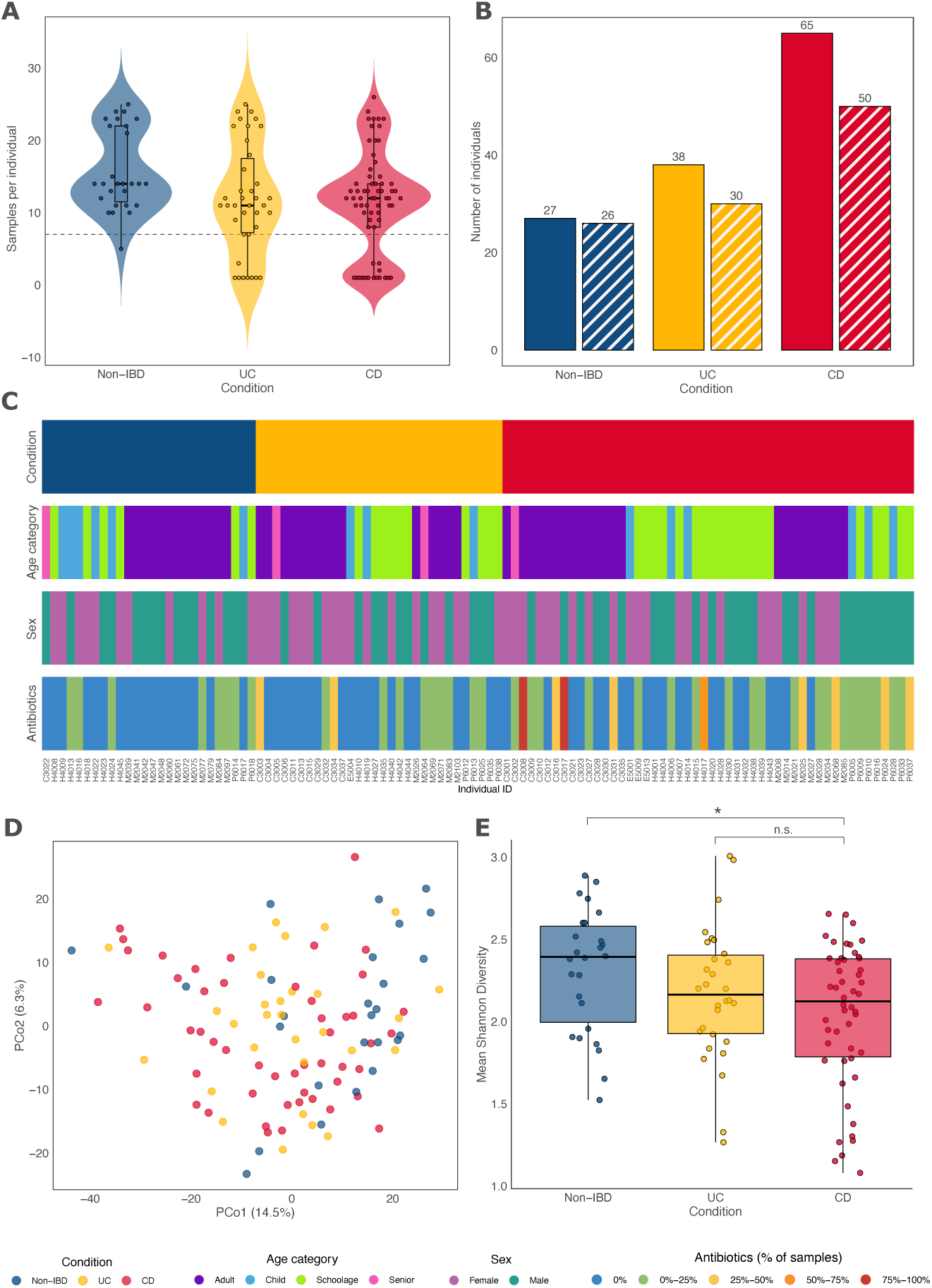
Overview and microbial diversity analysis of the discovery dataset. (**A**) Distribution of the number of longitudinal samples per individual. The dashed line indicates the threshold (*<* 7 samples) used to filter out individuals. (**B**) Number of individuals in the discovery dataset before (solid fill) and after (hatched fill) filtering using the threshold shown in A. (**C**) Visual representation of individu metadata, including condition, age category, sex, and the percentage of timepoints at which antibiotic use was reported. (**D**) Principal Coordinate Analysis (PCoA) ordination of Aitchison distances between individuals computed on mean CLR-transformed species abundances. (**E**) Box plots showing the distribution of mean Shannon diversity values within each condition. The asterisk represents statistical significance for a one-sided Mann-Whitney U test (*p* = 0.0073, testing the hypothesis that non-IBD diversity is greater than IBD [union of UC and CD individuals]). Differences between UC and CD were evaluated using a two-sided Mann-Whitney U test (*p* = 0.34). n.s., non-significant.

Among the remaining individuals, 26 were labeled as non-IBD and 80 as IBD (n = 30 UC and n = 50 CD) (Fig. 1B). Sex distribution was balanced, with 52 females and 54 males. Most of the individuals were adults (n = 51), followed by adolescents of school age (n = 36), children (n = 15), and seniors (n = 4). Only 11 individuals reported antibiotic use in more than 25% of their sampling visits, and none of these were non-IBD. Most individuals reported no antibiotic use (n = 58) or use in up to 25% of visits (n = 37) (Fig. 1C).

The dataset from Hall et al. (2017) was utilized for external validation, here termed the validation dataset. Specifically, it was used to evaluate the models trained on the discovery dataset and to assess the network-level features identified during model interpretation. This dataset comprises 259 longitudinal samples from 32 individuals. However, after applying the same filtering criteria from the discovery dataset, data from 25 adult individuals (n = 7 non-IBD, n = 9 UC, and n = 9 CD) remained (Suppl. Fig. 1A-C). Information regarding sex and antibiotic usage was unavailable for this dataset (Suppl. Fig. 1C).

We confirmed that, overall, within-individual Aitchison distances were lower than between-individual distances across samples for both datasets (Suppl. Fig. 2). Thus, we assumed that mean centered log-ratio (CLR)-transformed abundances across longitudinal samples could be used to represent each individual’s average gut microbiota composition. Using this metric, we then performed microbial diversity analyses at the individual level. Although the groups in the discovery dataset showed considerable overlap in the ordination of beta diversity (i.e., differences in community composition between individuals; see Methods, “Diversity analysis”), IBD individuals (UC and CD) exhibited a greater dispersion along the first principal coordinate (PCo1), which accounted for 14.5% of variance (Fig. 1D). In contrast, non-IBD individuals showed a notable spread along the second principal coordinate (PCo2), which accounted for 6.3% of variance (Fig. 1D). Furthermore, non-IBD individuals exhibited a significantly higher alpha diversity (i.e., the diversity of species within an individual’s microbiota, measured with the Shannon index) compared to those with IBD (Mann-Whitney U test, *p* = 0.0073), while no difference was observed between CD and UC (*p* = 0.34) (Fig. 1E). This reduction in alpha diversity in IBD relative to non-IBD is consistent with previous reports of dysbiosis in active disease (Lloyd-Price et al., 2019; Nishida et al., 2018).

The first principal coordinate (PCo1) in the ordination of beta diversity for the validation dataset accounted for 23.1% of variance while the second principal coordinate (PCo2) accounted for 9.3% (Suppl. Fig. 1D). Although condition overlap was also detected, non-IBD individuals were more concentrated on the left side of PCo1, whereas IBD individuals, particularly those with UC, occupied the right side. No significant differences in alpha diversity between conditions were detected in the validation dataset, which includes only 25 individuals (Suppl. Fig. 1E).

Stratified diversity analysis performed for the discovery dataset showed that alpha diversity in senior individuals (*>* 65 years) and in individuals with sustained antibiotic use differed from other groups (Suppl. Fig. 3A). However, no distinct clustering was observed in beta diversity for these same strata (Suppl. Fig. 3B). Although the number of individuals in these groups limits robust conclusions (seniors, n = 4; antibiotics in > 50% of the samples, n = 3), alterations in the gut microbiota associated with advanced age and antibiotic exposure have previously been reported (Yatsunenko et al., 2012; Palleja et al., 2018). This stratified diversity analysis was not performed for the validation dataset due to the lack of antibiotic usage and sex metadata, as well as age variation across individuals (Suppl. Fig. 1C).

### Individual-level microbial association networks preserve gut microbiome interindividual heterogeneity and condition overlap

We generated individual-level MANs by applying SparCC (Friedman and Alm, 2012) to each individual’s longitudinal samples independently (Fig. 2A). Similar approaches employing other correlation methods have been used to obtain individual-level brain networks (Gratton et al., 2018; Braga and Buckner, 2017; Finn et al., 2015). However, a key limitation of such strategies is that they may introduce spurious correlations driven by confounding temporal trends (Kodikara et al., 2022). To address this limitation, we applied linear detrending to the CLR-transformed data (see Methods, “Construction of individual-level microbial association networks”). By default, SparCC computes correlations for every pair of species present in the dataset, which generates fully connected networks that lack structural variation. To induce variation, the individual-level MANs were sparsified by eliminating edges with absolute weights below 0.3.

**Figure 2:**
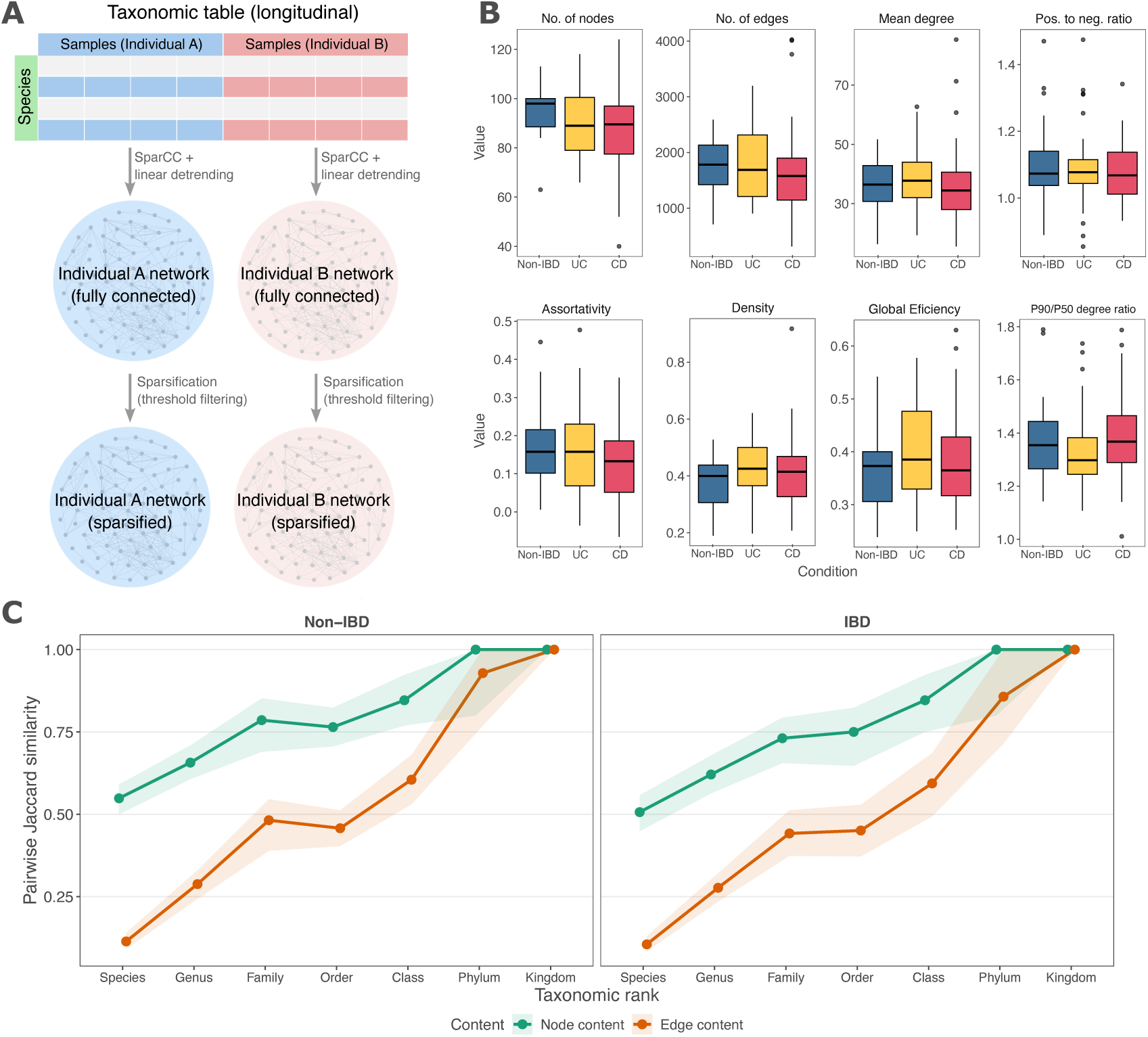
Construction of individual-level microbial association networks. (**A**) Schematic representation of the network construction methodology. A modified SparCC algorithm (incorporating linear detrending per species) was applied separately to each individual’s longitudinal sample set to construct individual-level microbial association networks (MANs). In these networks, nodes represent bacterial species and edges represent SparCC weighted associations. Individual-level MANs were subsequently sparsified by filtering edges using absolute association weight thresholds (0.3, 0.4, or 0.5). For simplicity, the workflow is illustrated for only two individuals. (**B**) Box plots showing the distributions of computed network metrics for individual-level MANs across different conditions. These metrics were calculated specifically for MANs sparsified using an absolute weight threshold of 0.3. No significant differences were detected by comparing conditions within each metric (two-sided Mann Whitney U test with Benjamini-Hochberg correction, FDR *>* 0.05 for all comparisons). (**C**) Pairwise Jaccard similarities for node and edge content. Similarities were calculated as the taxonomic resolution of the nodes was progressively aggregated from species to higher taxonomic levels, separately for non-IBD and IBD networks. Points and connecting lines represent the median Jaccard similarity across individual pairs, while the shaded ribbons denote the interquartile range (IQR; 25th to 75th percentiles). microbiota Backhed et al. (2005).

An analysis of network metrics revealed substantial heterogeneity between individuals, as evidenced by the variability observed in the boxplots in Figure 2B. However, when comparing conditions, no significant differences were observed between the non-IBD and IBD groups, nor between the non-IBD group and specific IBD subtypes (UC and CD) (Mann-Whitney U test, FDR *>* 0.05 for all comparisons). These results indicate that, despite individual variations, there is a degree of structural overlap in the networks across conditions.

Additionally, we analyzed the similarity between MANs regarding their node and edge sets by calculating the Jaccard similarity index. We observed that similarity between node sets was consistently higher than that observed for edge sets, and this pattern occurred regardless of the clinical condition (non-IBD or IBD). However, as the nodes’ taxonomic resolution shifted to broader categories, we noted that the networks became progressively more similar to one another (Fig. 2C). That is, the node and edge heterogeneity of the MANs was attenuated by taxonomic generalization, indicative of lower taxon richness at broader taxonomic levels (such as phyla) compared to finer levels (such as species), a phenomenon commonly observed in the human gut

### Traditional models outperform GNNs on small, imbalanced metagenomic data

We compared the classification performance of GNNs trained on individual-level MANs against traditional models trained exclusively on tabular abundance data from the same discovery dataset. Specifically, models were evaluated on a binary classification task to distinguish between non-IBD and IBD individuals. Given the limited sample size (n*_individuals_* = 106) and the notable class imbalance (approximately 1:3 non-IBD to IBD ratio), we implemented Monte Carlo-like experiments consisting each of 50 independent, randomized train-test splits from the discovery dataset, followed by an external validation on the validation dataset. Each individual experiment evaluated a model trained on a specific combination of node attributes and network sparsification threshold.

To prepare these individual-level MANs for graph-based machine learning, we enriched the nodes with two distinct categories of attributes. First, taxonomic attributes were obtained by constructing a one-hot encoded vector representing the complete taxonomic lineage of each species, from genus to phylum. Second, abundance attributes were structured as a two-element vector containing the species-specific mean CLR-transformed abundance within that individual, alongside its associated log-transformed variance (Fig. 3A). To use network structure information only, we also trained models with dummy attributes, assigning the same constant value to every node (see Methods, “Node attributes”).

**Figure 3:**
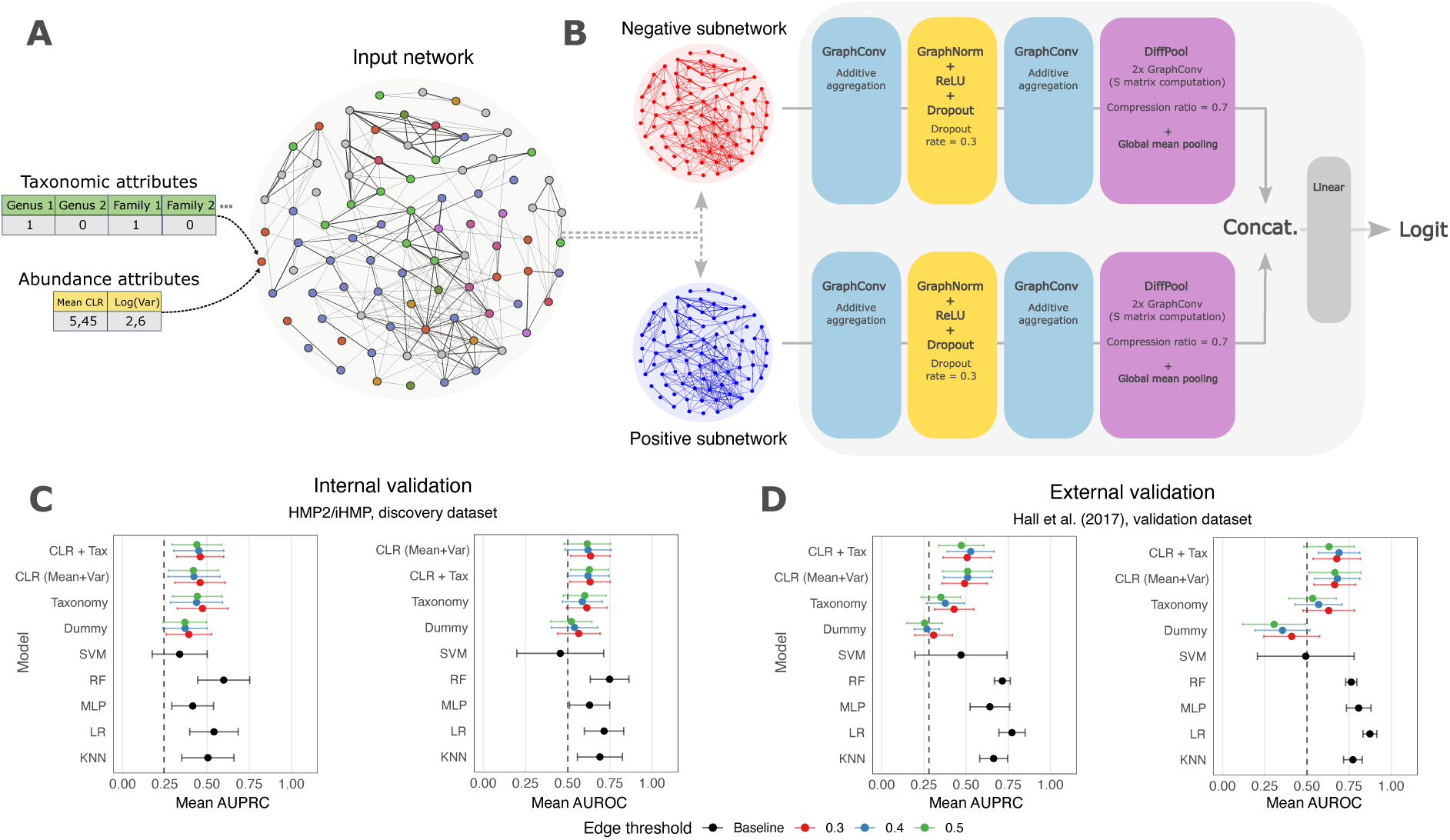
Classification performance of GNNs and traditional machine learning models. (**A**) Schematic of the input network preparation for GNN training. Sparsified individual-level MANs were enriched with biological features via node attribute vectors. Taxonomic attributes were generated by one-hot encoding each node’s taxonomy into a vector representing all possible taxa (from genus to phylum) observed in the discovery dataset. Abundance attribute vectors consisted of individual-specific mean CLR-transformed abundances and their associated logtransformed variances. (**B**) The GNN architecture featured two identical processing streams corresponding to a positive and a negative subnetwork derived from the input network. The positive subnetwork comprised positive edges only, while the negative subnetwork contained negative edges with their weights converted to absolute values. Graph representation vectors from both streams were concatenated before passing through the final linear layer for classification (Zhang et al., 2024). (**C**) Mean AUPRC and AUROC values for classification experiments during internal validation on the discovery dataset (HMP2/iHMP; Lloyd-Price et al. (2019)). Dashed lines indicate the baseline for each metric (minority class proportion for AUPRC; 0.5 for AUROC). Points represent mean values computed over 50 independent Monte Carlo-like experiments, with error bars indicating standard deviations. GNNs trained on different node attribute combinations are ordered by their mean performance across the three edge thresholds, with the best-performing combination at the top. Traditional machine learning models are shown below for comparison. Traditional machine learning benchmarks include Support Vector Machine (SVM), Random Forest (RF), Multi-Layer Perceptron (MLP), Logistic Regression (LR), and *K*-Nearest Neighbors (KNN). (**D**) Mean AUPRC and AUROC values for classification experiments performed during external validation on the dataset from Hall et al. (2017).

The GNN architecture was specifically designed to handle the presence of both positive and negative edges through a dual-stream framework (Zhang et al., 2024). For each network, the structure was divided into two identical parallel streams: one processing a subnetwork composed entirely of the positive edges, and the other processing a subnetwork of only the negative edges with their weights transformed to absolute values (Fig. 3B). This explicit separation was implemented to prevent opposing edge weight signals from canceling each other out during message passing, ensuring that both positive and negative signals were independently retained. At the final stage of the architecture, the graph representation vectors generated by both streams were concatenated and passed through a final linear layer to output logits (Fig. 3B).

After performing the Monte Carlo-like experiments, we observed that most of the traditional models trained on tabular abundance data exhibited higher average classification performance than GNNs, while most models in both approaches displayed similar performance instability during internal validation (Fig. 3C). Among the traditional models, Random Forests (RFs) achieved the best performances during internal validation, with the highest average AUPRC (0.599 *±* 0.154, mean *±* SD) and AUROC (0.748 *±* 0.115) scores relative to other models (Fig. 3C), while Logistic Regression (LR) models performed the best during external validation (AUPRC, 0.772 *±* 0.077; AUROC, 0.872 *±* 0.041) (Fig. 3D). In contrast, Support Vector Machines (SVMs) exhibited the lowest performance and stability, with high variance of the AUROC metric during internal validation (0.456 *±* 0.258) (Fig. 3C) and both AUPRC (0.470 *±* 0.273) and AUROC (0.493 *±* 0.287) during external validation (Fig. 3D).

When comparing different combinations of node attributes during internal validation, we observed that omitting biological information from the MANs resulted in lower classification performances for the GNNs. Because each configuration was evaluated at three sparsification thresholds (0.3, 0.4, and 0.5), we report the ranges of means and standard deviations across thresholds and, where applicable, attribute combinations. Models with dummy attributes reached mean AUPRC of 0.369–0.392 (SD 0.129–0.133) and mean AUROC of 0.522–0.565 (SD 0.121– 0.137), compared with mean AUPRC of 0.419–0.473 (SD 0.139–0.154) and mean AUROC of 0.587–0.635 (SD 0.112–0.138) for configurations with biological attributes (Fig. 3C). This gap was also present during external validation between dummy attributes (mean AUPRC 0.254– 0.308, SD 0.075–0.111; mean AUROC 0.305–0.410, SD 0.162–0.186) and configurations with biological attributes (mean AUPRC 0.350–0.527, SD 0.111–0.148; mean AUROC 0.534–0.691, SD 0.123–0.154). Attribute combinations that included mean CLR-transformed abundances ranked the highest for both AUPRC and AUROC in the internal and external validations, suggesting a meaningful contribution of species abundance profiles in discriminating between conditions, even in a graph-based setting (Fig. 3C-D).

### Bacteroidales rank as the most important nodes for GNN classification

While traditional models provided strong predictive performance, GNNs enabled us to investigate network-level features relevant to condition discrimination. Unlike standard feature attribution methods that evaluate variables independently, this approach incorporates topological context to capture complex, higher-order interactions. To identify these relevant network-level features, we selected 30 distinct, well-fitting GNN model instances from 50 Monte Carlo-like runs (see Methods, “Interpretation of graph neural networks via Integrated Gradients”). These models were trained on individual-level MANs containing taxonomic, and abundance node attributes, with network sparsification performed using an absolute association weight threshold of 0.3.

We applied the Integrated Gradients method (Sundararajan et al., 2017) to the selected models, extracting importance scores for every node attribute and edge across individual-level MANs (see Methods, “Interpretation of graph neural networks via Integrated Gradients”). For each model, attribute scores were summed per node to obtain node-level importance, both as signed values (net contribution toward a given class) and as absolute values (cumulative contribution irrespective of direction). These scores were then averaged across models and max-normalized within each graph to derive final signed and absolute node importances. Nodes were ranked by their mean absolute importance across all discovery MANs, weighted by node prevalence (the proportion of MANs in which each node was present), revealing the top 20 nodes prioritized by the GNNs (Fig. 4).

**Figure 4:**
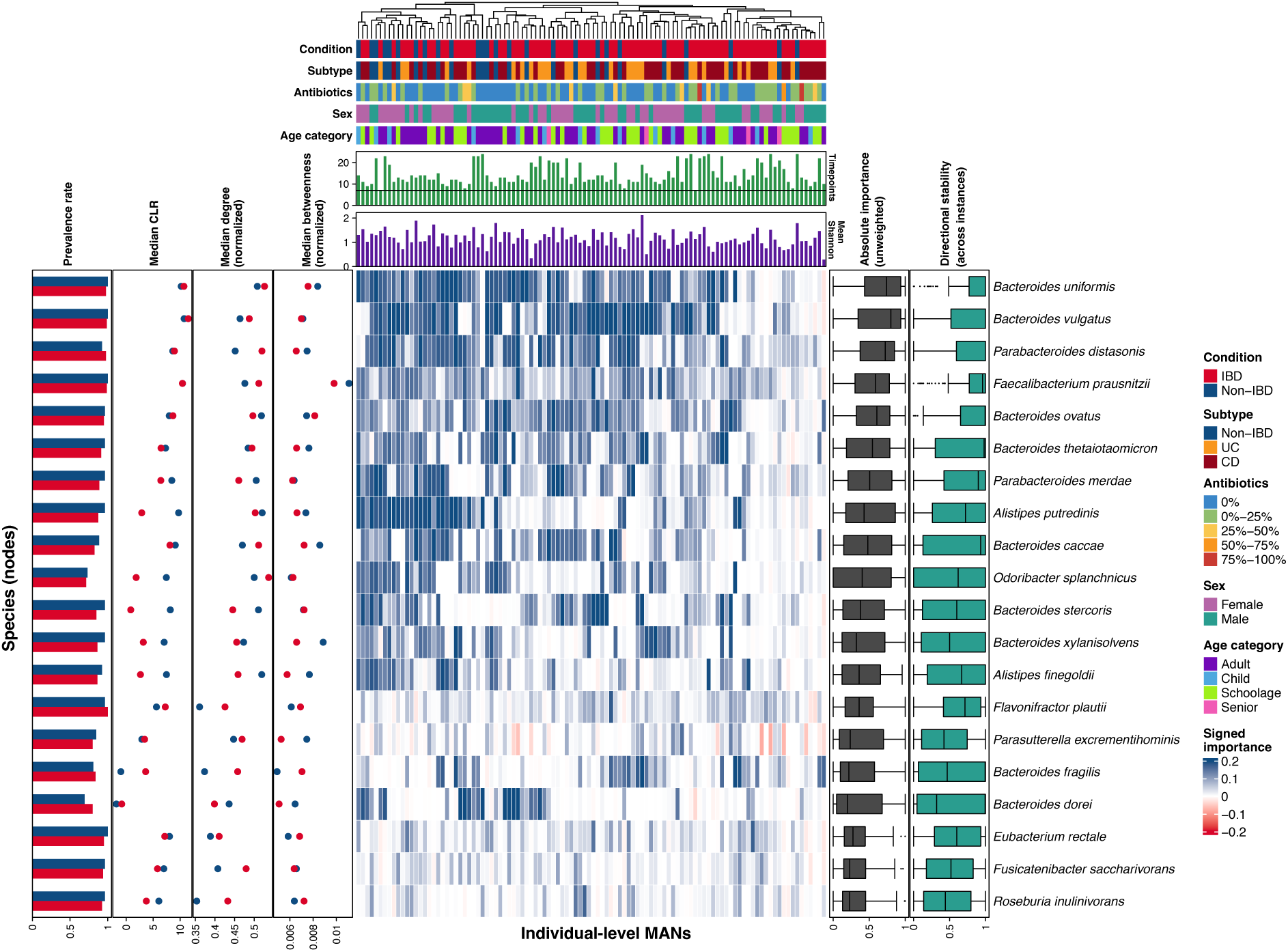
Top 20 nodes prioritized by GNNs. Heatmap displaying the top 20 prioritized nodes based on prevalence-weighted mean absolute importance computed across MANs. Rows represent species (nodes) and columns correspond to individual-level MANs. The column order reflects hierarchical clustering using Euclidean distance and complete linkage. The central heatmap color intensity represents signed importance (red = negative scores toward IBD; blue = positive scores toward non-IBD). Top annotations include individual-level clinical metadata, number of longitudinal samples (timepoints), and mean Shannon diversity. The black horizontal line on the timepoints barplot indicates the minimum inclusion threshold of 7 longitudinal samples. Left annotations show species-level prevalence rate, median CLR, median normalized degree, and median normalized betweenness per condition. Right annotations show distributions of node absolute importance and signed importance directional stability (i.e., the consistency of signs across distinct model instances). Statistical differences between IBD and non-IBD individuals were evaluated for each left annotation metric. Fisher’s exact test was applied to individual presence counts to compare prevalence, and two-sided Mann-Whitney U tests were applied to CLR-transformed abundance and normalized centrality metrics. *p*-values were adjusted using the Benjamini-Hochberg procedure, and no comparison was significant (FDR > 0.05 for all).

Among the top 20 nodes, signed importance scores followed two main patterns across MANs: consistently positive, or neutral-to-positive (Fig. 4, central heatmap). Nodes with consistently positive scores were stably associated with the non-IBD class, whereas neutral-to-positive nodes showed more near-zero signed scores, reflecting lower directional stability across model instances (Fig. 4, right annotations). None of the top 20 nodes showed consistently negative scores, suggesting that the evidence used by the GNNs was predominantly non-IBD-directed, and that IBD MANs were distinguished by weaker positive contributions from these taxa rather than by nodes pointing toward IBD.

Absolute node importance was associated with node centrality and species abundance (Fig. 5). Weighted mean absolute importance correlated moderately with median node degree (Spearman *ρ* = 0.55, *p* = 2 *×* 10^−14^) (Fig. 5A) and median betweenness (*ρ* = 0.68, *p <* 2 *×* 10^−16^) (Fig. 5B), and strongly with CLR-transformed abundance (*ρ* = 0.9, *p <* 2 *×* 10^−16^) (Fig. 5D). Median degree and median betweenness were also moderately correlated (*ρ* = 0.52, *p* = 8.2 *×* 10^−13^) (Fig. 5C). Furthermore, decomposing node importance by attribute type showed that abundance and taxonomic attributes contributed comparably, with abundance accounting for a slightly larger share among the top 20 nodes (53–65%) than across all nodes (48%) (Suppl. Fig. 4). Nevertheless, the top 20 nodes did not differ significantly between conditions in prevalence, abundance, degree, or betweenness (FDR > 0.05 for all comparisons), and they did not overlap with the features prioritized by RF models, all of which were differentially abundant between conditions (Suppl. Fig. 5). Together, these results indicate that the GNNs relied mainly on dominant (i.e., highly abundant) taxa that were also highly connected and occupied bridging positions in the MANs, but that do not distinguish conditions in univariate comparisons. Graph-based and traditional models therefore appear to capture complementary aspects of the data.

**Figure 5:**
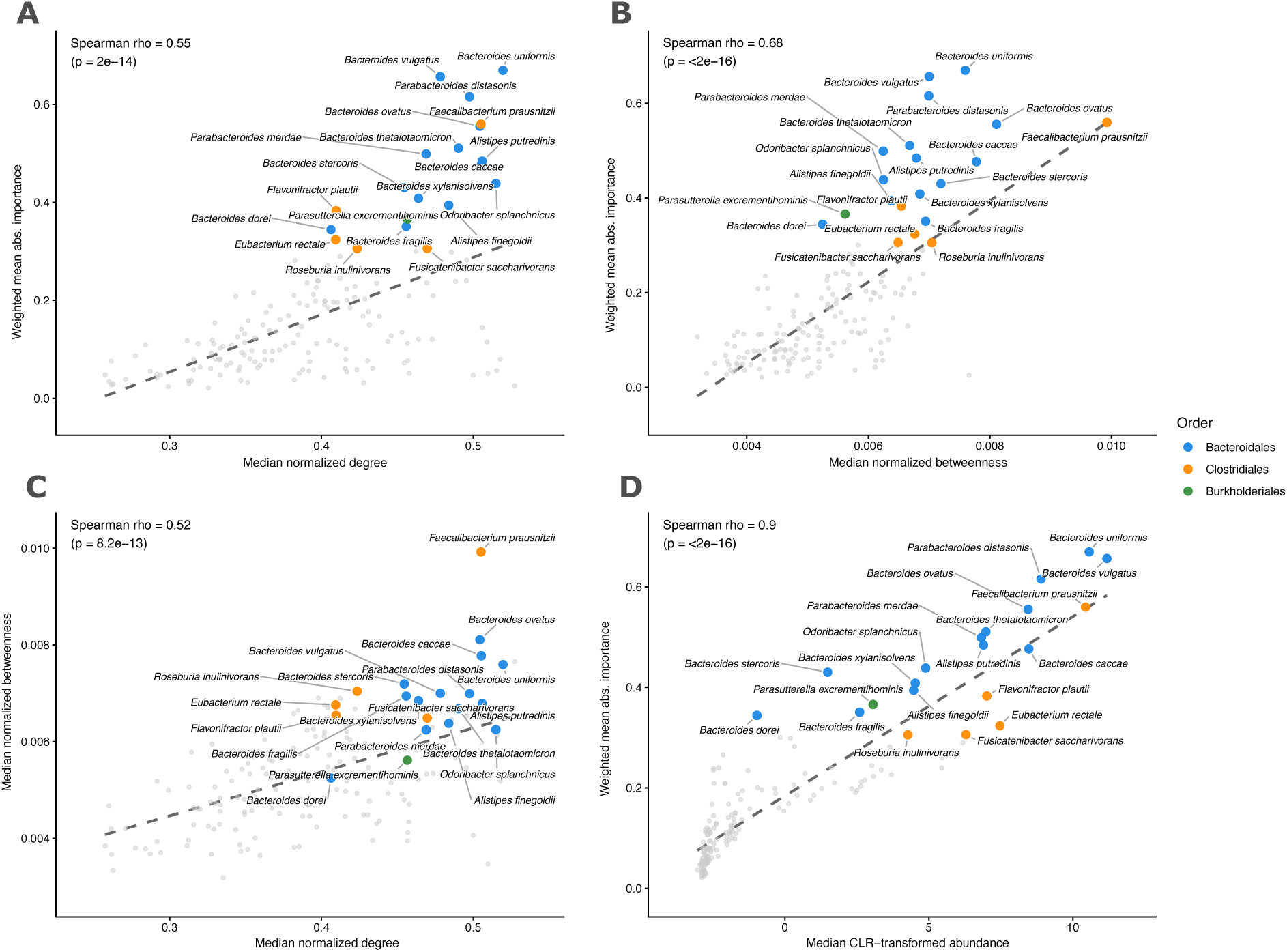
Node importance in relation to network topology and abundance. Each point represents a species (node), with node-level metrics summarized across all individual-level MANs. The top 20 prioritized nodes are highlighted and colored by taxonomic order. (**A**) Weighted mean absolute node importance versus median normalized degree. (**B**) Weighted mean absolute node importance versus median normalized betweenness centrality. (**C**) Median normalized betweenness centrality versus median normalized degree. (**D**) Weighted mean absolute node importance versus median CLR-transformed abundance. Dashed lines show linear fits. Spearman’s correlation coefficients (*ρ*) and p-values are given in each panel.

The top 20 nodes were dominated by Bacteroidales, which accounted for 14 species, including the three highest-ranked nodes (*Bacteroides uniformis*, *B. vulgatus*, and *Parabacteroides distasonis*) (Fig. 4). Bacteroidales are highly prevalent obligate anaerobes that degrade dietary polysaccharides and produce short-chain fatty acids (SCFA), and their depletion has been reported in treatment-naive IBD (Gevers et al., 2014), although abundance-level associations vary across studies and with disease severity (Zhou and Zhi, 2016; Nomura et al., 2021; Mills et al., 2022). The remaining top nodes were mostly Clostridiales, including the butyrate producers *Faecalibacterium prausnitzii*, *Eubacterium rectale*, and *Roseburia inulinivorans* (Louis and Flint, 2009). Among these, *F. prausnitzii*, a well-documented anti-inflammatory species whose depletion is associated with gut dysbiosis and colitis (Sokol et al., 2008, 2009; Quévrain et al., 2016), ranked fourth (Fig. 4) and held the highest median betweenness among the top 20 nodes (Fig 5B-C).

### The predictive signal of individual-level MANs is concentrated in associations among Bacteroidales species

Node-level scores aggregate the contributions of all node attributes into a single value and therefore cannot resolve which of a node’s associations carried the signal. Edge scores, indexed by node pairs, provide the importance of specific associations. Therefore, to capture the most important associations, we ranked the edges using prevalence-weighted mean absolute importance and selected the top 100.

The largest connected component of the resulting importance metagraph was dominated by a densely connected module of Bacteroidales species (17 of 22 nodes; Fig. 6B), whose edges carried positive mean signed importance (i.e., evidence toward the non-IBD class) (Fig. 6A). The strongest edges linked *B. uniformis*, *B. vulgatus*, and *P. distasonis*, the three top prioritized nodes (Fig. 4), and *F. prausnitzii* and *E. rectale* were the only Clostridiales connected to the module. A small branch of three Proteobacteria species was attached to the module by a single low-importance edge. At the genus level, most nodes belonged to *Bacteroides* (10 of 22; Fig. 6B). Together, node- and edge-level interpretation converged on the same Bacteroidales species, which ranked as the most important nodes and formed the strongest associations of the metagraph. To assess whether this metagraph carried the predictive signal of the MANs, we trained models on the discovery dataset using three graph configurations: the whole graph (WG), the whole graph with metagraph nodes and their incident edges removed (WG - M), and metagraph-only graphs (M only), in which only metagraph nodes and the edges among them were retained. To separate the contribution of associations from that of node attributes, each configuration was also trained with all edges removed, keeping all nodes and node attributes. Each trained model was then evaluated on the validation dataset (Fig. 7). Results for all training and evaluation combinations, including AUROC, are provided in Supplementary Table 4.

**Figure 6:**
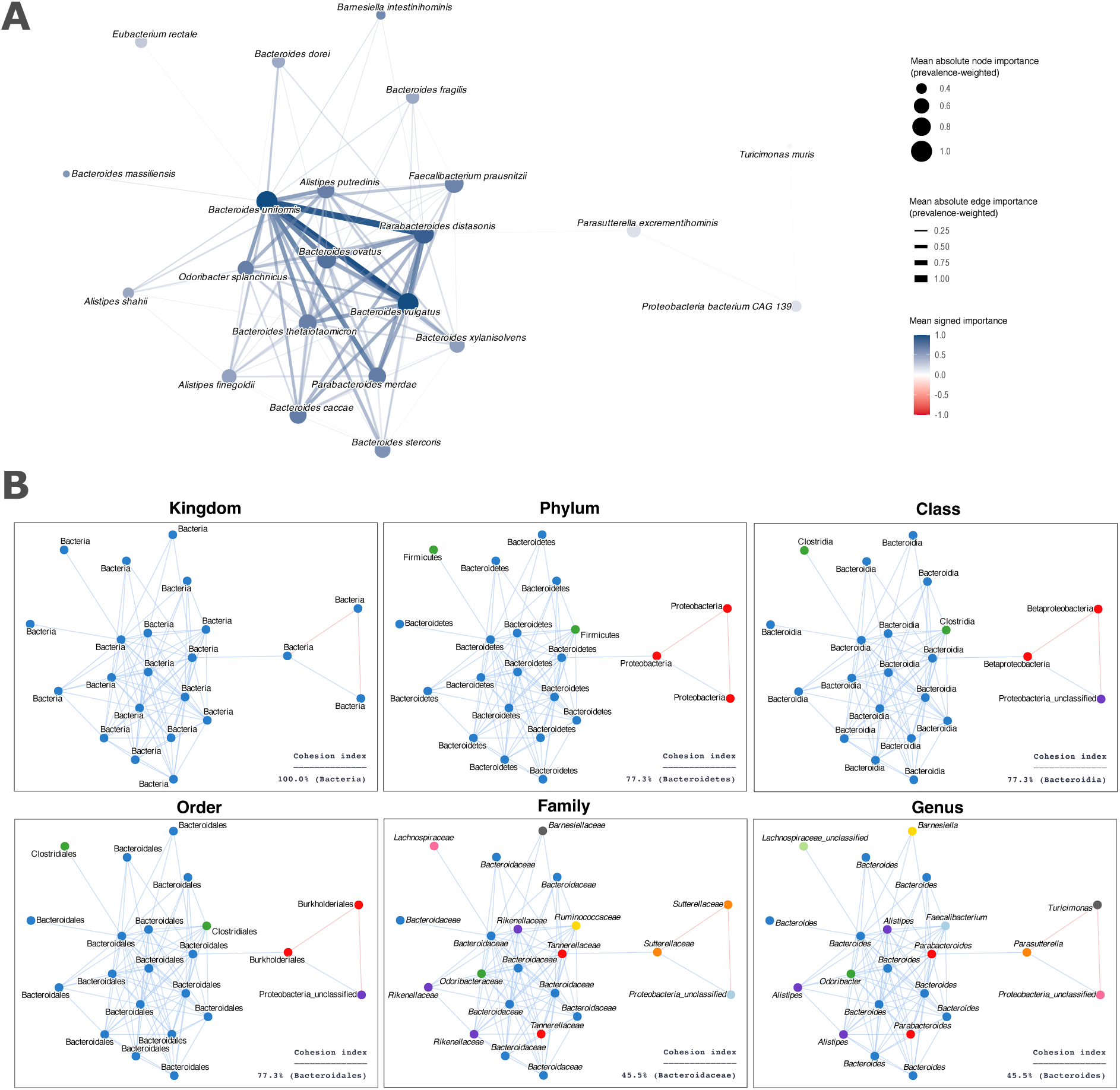
Importance metagraph formed by the top 100 prioritized edges. (**A**) The largest connected component formed by the 100 edges with the highest prevalence-weighted mean absolute importance across MANs. Edge thickness and node size represent prevalence-weighted mean absolute importance. Colors represent mean signed importance (red, IBD-directed; blue, non-IBD-directed). Values were scaled for visual clarity. (**B**) Taxonomic composition of the metagraph from kingdom to genus. Node colors designate distinct taxa within each panel, and edge colors indicate the sign of mean signed importance. The cohesion index is the proportion of nodes belonging to the most common taxon at each level.

**Figure 7:**
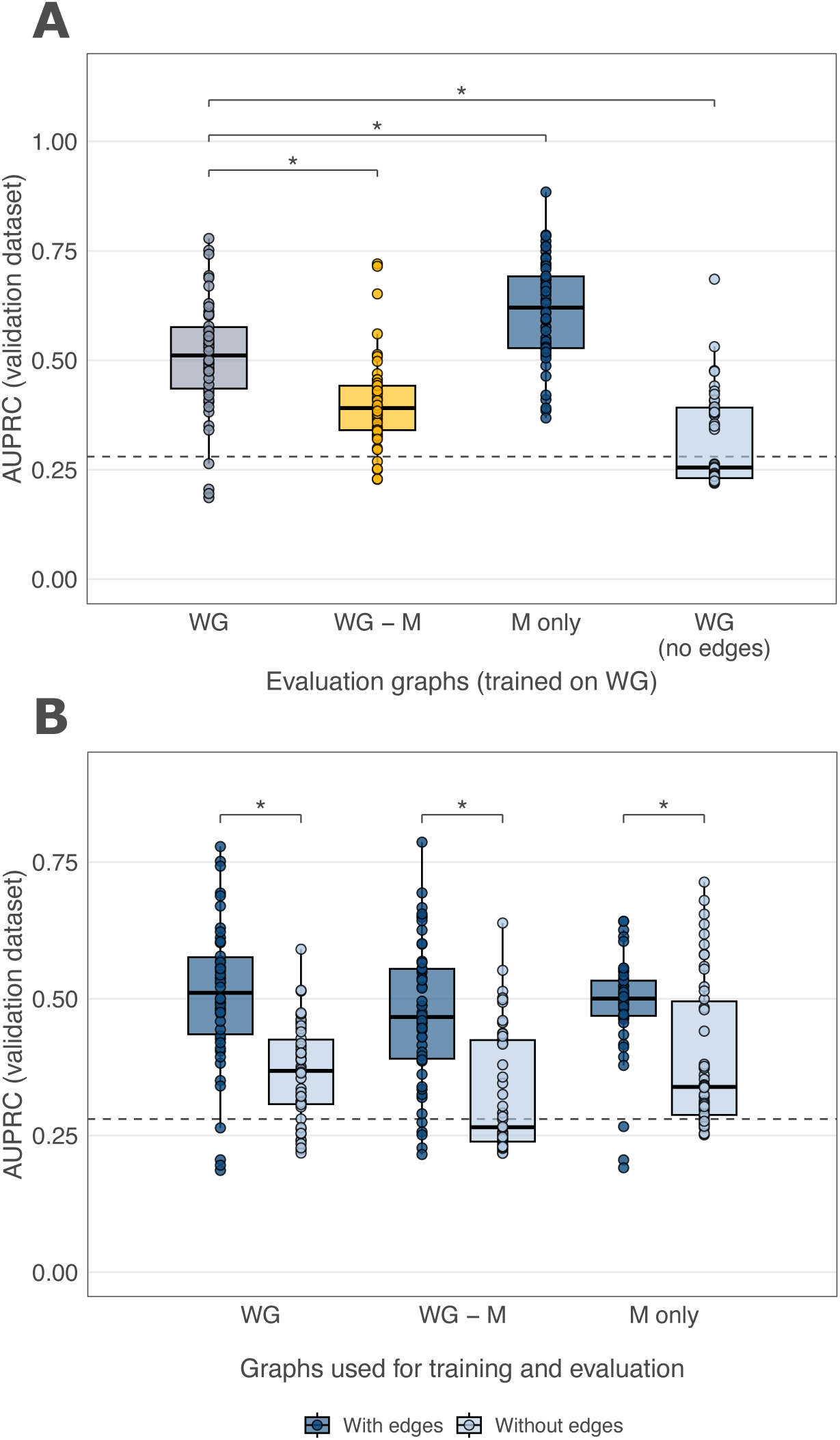
GNN performance after removing metagraph nodes or edges. (**A**) AUPRC of models trained on the whole graph (WG) and evaluated on the validation dataset using whole graphs (WG), graphs with metagraph nodes removed (WG - M), metagraph-only graphs (M only), and whole graphs without edges. (**B**) AUPRC of models trained and evaluated on the same graph configuration, with and without edges. Graphs without edges keep all nodes and node attributes. Each point represents one of 50 Monte Carlo-like runs. The dashed line represents the baseline AUPRC for random classification (proportion of non-IBD individuals in the validation dataset; 0.28). Significance was assessed using two-sided Wilcoxon signed-rank tests paired by model run, with Benjamini–Hochberg correction across all comparisons (*, FDR < 0.05). Results for all training and evaluation combinations including AUROC values can be found in Supplementary Table 4.

Models trained on WG performed best when evaluated on M-only graphs (AUPRC, 0.614 *±* 0.117 vs. 0.507 *±* 0.131 on WG; FDR = 1.0 *×* 10^−4^) and dropped when metagraph nodes were removed (AUPRC, 0.403 *±* 0.103; FDR = 2.5 *×* 10^−7^) (Fig. 7A). Removing all edges reduced performance to near chance (AUPRC, 0.316 *±* 0.103; FDR = 4.0 *×* 10^−7^). When models were trained and evaluated on the same configuration, removing edges lowered performance for WG (AUPRC, 0.507 *±* 0.131 vs. 0.364 *±* 0.087; FDR = 1.1 *×* 10^−6^), WG - M (0.471 *±* 0.130 vs. 0.324 *±* 0.110; FDR = 1.4 *×* 10^−5^), and M only (0.491 *±* 0.088 vs. 0.391 *±* 0.133; FDR = 1.3 *×* 10^−4^) (Fig. 7B). These results indicate that the predictive signal of the MANs depends on associations rather than on node attributes alone, that associations among metagraph species are sufficient to retain it, and that this signal generalizes to an independent cohort.

To characterize the associations underlying this signal, we compared SparCC weights among the 17 Bacteroidales species of the metagraph between non-IBD and IBD individuals. These associations had positive SparCC weights in nearly all individuals of both conditions, and their mean weight per individual was higher in non-IBD than in IBD individuals in both the discovery (Mann-Whitney U test, *p* = 0.042) and the validation dataset (*p* = 0.007) (Suppl. Fig. 6A). This difference was not driven by a few pairs of species. Instead, most pairwise associations had higher mean weights in non-IBD individuals in both cohorts (108 of 136 pairs in the discovery and 80 of 87 in the validation dataset), with *Bacteroides fragilis* as the main exception (Suppl. Fig. 6B). The predictive signal of the metagraph thus appears to reflect a cohesive module of co-abundant Bacteroidales, with weaker associations in IBD individuals of both cohorts.

## Discussion

In this study, we combined metagenomics and graph-based machine learning to ask what can be learned from interpreting models trained on individual-level microbial association networks (MANs) for IBD classification. We based our approach on the view that microbiotas are complex ecological systems, in which microorganisms interact directly and indirectly, cooperating and competing for resources to thrive within their niches (Faust and Raes, 2012), and that MANs can serve as a proxy for that organization. Representing each individual as a network allowed us to move beyond asking which taxa differ between conditions and instead investigate which associations between taxa carry the signal. Although traditional models trained on abundance tables achieved higher overall predictive performance, the features they prioritized differed from those of the graph neural networks (GNNs). Random Forests ranked species that were differentially abundant between conditions, none of which were among the nodes prioritized by GNN-based interpretation (Suppl. Fig. 5). Instead, the GNNs relied on a densely connected module of co-abundant Bacteroidales species that carried most of the predictive signal in an independent cohort.

Inferring MANs that capture associations between taxa (Faust and Raes, 2012) typically requires many observations, because standard inference metrics such as correlation coefficients estimate network edges directly from the covariation of abundances across samples (Weiss et al., 2016). Consequently, studies frequently construct MANs at the cohort level to generate a single, global association network (Faust et al., 2012; Zhang et al., 2014; Chen et al., 2020). However, cohort-level inference can mask real heterogeneity between the conditions of interest, as in Simpson’s paradox (Simpson, 1951). Inferring one network per condition (Lam and Ye, 2022; Geese et al., 2025) offers a partial workaround, but the resulting associations still cannot capture the inter-individual heterogeneity inherent to the gut microbiome, and they yield one network per group rather than one per individual, which precludes their use as input samples for machine learning.

Sample-specific network approaches, such as LIONESS (Kuijjer et al., 2019), SWEET (Chen et al., 2023), and SSN (Liu et al., 2016), instead derive one network per individual by estimating each individual’s contribution to a global network. They can be applied to cross-sectional data, but every resulting network inherits the topology of the global network and individuals differ only in edge weights. Those weights are typically not bounded to a fixed interval (De Marzio et al., 2023), which complicates comparisons across networks. Inferring each network independently instead preserves differences in both node and edge content between individuals (Fig. 2C).

Graph-based machine learning has only recently been applied to microbiome data. Beyond related metagenomic tasks such as contig binning (Lamurias et al., 2022; Xue et al., 2022; Wei and Liu, 2025), which address a different problem, most methods for host phenotype prediction build the graph from something other than inferred microbial associations. Shtossel et al. (2023), for instance, represented each sample as the taxonomic tree with abundances as node attributes, and applied graph and convolutional neural networks to 16S microbiome data. Syama et al. (2023) proposed a disease-agnostic framework to predict healthy and diseased states in which each metagenomic sample was a node and edges connected samples according to patient similarity. Methods that do use inferred MANs have generally estimated them at the cohort or class level in order to classify networks representing diseased versus healthy populations (Xu et al., 2022; Pan et al., 2024). By contrast, our approach infers a signed MAN for each individual, which allows graph classification at the individual level and interpretation in terms of specific associations, at the cost of using a limited number of longitudinal samples.

We observed that, overall, traditional models achieved higher classification performance than GNNs, with Random Forests (RFs) and Logistic Regressions (LRs) emerging as the top performers, showing that microbial abundance tables remain highly effective for individual classification. This agrees with previous comparisons of traditional machine learning and deep learning on taxonomic abundance data (Li et al., 2025; Topçuoğlu et al., 2020; Pasolli et al., 2016). The advantage of RFs and LRs likely reflects their suitability for small datasets, since both have few parameters to tune, RFs are inherently regularized through bagging, and our LRs were stabilized by *L*_2_ regularization, which manages multicollinearity among correlated taxa. In contrast, graph-based and other complex frameworks are highly parametrized, sensitive to architecture and hyperparameters, and generally data-hungry (Hu et al., 2021), which makes them harder to train and generalize on small and imbalanced datasets such as the HMP2/iHMP cohort aggregated to individuals (*n_individuals_* = 106; 1:3 non-IBD to IBD ratio).

Beyond predictive performance, GNNs allowed us to ask where in the network the discriminative signal lies. Global network metrics did not differ between IBD and non-IBD individuals (Fig. 2B), and models relying only on topology and edge weights performed worse than models trained on MANs containing node attributes (Fig. 3C-D). The nodes prioritized by interpretation did not differ between conditions either, in prevalence, abundance, or centrality. What did differ were the associations among them, which were weaker in IBD individuals in both cohorts (Suppl. Fig. 6). This suggests that GNN-guided analysis can recover differences that neither global topological summaries nor taxon-level comparisons capture.

The importance metagraph was dominated by a dense module of Bacteroidales species whose edges were directed toward the non-IBD class. Bacteroidales are obligate anaerobes that degrade dietary polysaccharides and produce short-chain fatty acids (SCFA), and together with the host immune system they are thought to restrain Enterobacteriaceae through colonization resistance (Spees et al., 2013). Some also cooperate directly, as in the cross-feeding between *B. ovatus* and *B. vulgatus* (Rakoff-Nahoum et al., 2016), both of which are among the prioritized species. A densely connected module of co-abundant Bacteroidales may therefore reflect a cohesive anaerobic community that is perturbed in active IBD, since the rise in oxygen levels during inflammation is thought to alter the anaerobic environment of the gut (Lupp et al., 2007; Rigottier-Gois, 2013). In the HMP2/iHMP cohort, the original authors (Lloyd-Price et al., 2019) reported that disease activity is marked by a loss of obligate anaerobes in favor of facultative anaerobes, though they did not highlight Bacteroidales among the taxa characterizing dysbiotic samples. We also found no abundance differences among these species, yet their associations were consistently weaker in IBD individuals of both cohorts. Co-abundance relationships have been shown to carry disease-specific information in larger IBD cohorts (Chen et al., 2020), and our results indicate that such information can be present in species whose abundances do not differ between conditions.

The study most closely related to ours is that of Ivanov et al. (2026), who trained graph-level classifiers on one graph per sample and used Integrated Gradients to derive node and edge importances in a food allergy microbiome dataset, outperforming earlier image- and graph-based deep learning methods and prioritizing taxa that largely agreed with the allergy literature. Their graphs were fully connected with uniform edge weights, which avoids prior assumptions but means that extracted edge importances reflect only what the model learned and not estimated associations between taxa. Additionally, their models were evaluated by cross-validation within a single dataset and compared with other deep learning methods, which leaves open whether the taxa and taxon pairs they identified generalize to independent cohorts, and how their performance compares with that of traditional models, which have shown high predictive performance on microbiome abundance data (Li et al., 2025; Topçuoğlu et al., 2020; Pasolli et al., 2016).

Several aspects of this study point to future work. First, both cohorts are small, particularly the validation dataset (25 adults, 7 of them non-IBD), and the class imbalance contributed to the variance across runs. Individual-level MANs were inferred from a limited number of longitudinal samples per individual, which also varied in number and spacing across individuals, and, because SparCC estimates pairwise correlations, edges represent statistical associations rather than direct ecological interactions. Inference methods that target direct associations, such as SPIEC-EASI (Kurtz et al., 2015), could further refine these networks. Similarly, species-level nodes may aggregate functionally distinct strains, which strain-resolved data could disentangle. Although importance scores were consistent across model instances, extending the analysis to other GNN architectures and attribution methods would establish how general the identified module is. Finally, all analyses are associative, so they may be influenced by factors such as antibiotic exposure, age, treatment, and disease activity, and longitudinal designs spanning disease onset will be needed to determine whether these network changes precede or follow disease.

Overall, interpreting GNNs trained on individual-level MANs revealed association-level features of the IBD gut microbiome that are not accessible to models trained on abundance tables alone. Although traditional models achieved higher predictive performance, GNN-based interpretation prioritized a densely connected module of co-abundant Bacteroidales species that, in contrast to the differentially abundant species prioritized by Random Forests, did not differ in abundance between conditions. Ablation experiments showed that this module carried most of the predictive signal in an independent cohort, and that removing its associations reduced performance to near chance. Together, these results complement abundance-based analyses and suggest that the organization of microbial associations, and not only the abundance of individual taxa, differs between IBD and non-IBD individuals.

## Methods

### Discovery and validation datasets

For individual-level MAN construction, model training, and interpretation, we used a metagenomic dataset of human fecal samples from individuals with and without IBD, generated by the HMP2/iHMP project (Lloyd-Price et al., 2019). The species-level taxonomic profiles and harmonized metadata were obtained from the curatedMetagenomicData R package (v3.14.0) (Pasolli et al., 2017), which provides uniformly processed MetaPhlAn 3 profiles (Beghini et al., 2021). Species abundances were retrieved as estimated read counts, as provided by the package, instead of relative abundances.

Following the age categories defined by Pasolli et al. (2017), individuals were grouped as newborn (*<* 1 year), child (1 *≤* age *<* 12), school age (12 *≤* age *<* 19), adult (19 *≤* age *≤* 65), or senior (*>* 65 years). We also defined an individual-level metric representing the proportion of longitudinal sampling timepoints at which antibiotic use was reported.

Before filtering, the discovery dataset comprised 1,627 longitudinal fecal samples from 130 individuals in the United States (64 females, 66 males; mean age = 27.5 years, SD = 17.35), including 103 individuals with IBD (65 with Crohn’s disease and 38 with ulcerative colitis) and 27 non-IBD controls.

To assess the generalizability of the models and of the network-level features identified during interpretation, we used an independent cohort from Hall et al. (2017), obtained from the same package. Before filtering, this validation dataset comprised 259 samples from 32 individuals, including 20 with IBD (9 with Crohn’s disease and 11 with ulcerative colitis) and 12 non-IBD controls. All individuals were adults, but numeric age and sex metadata were unavailable. Filtering, normalization, and network inference were applied to each dataset independently, using identical parameters and criteria.

### Diversity analysis

For microbial diversity analysis of the discovery and validation datasets, abundance tables were transformed using the centered log-ratio (CLR) transformation with a pseudocount of 1 to account for compositionality. CLR-transformed abundances were then aggregated by individual, calculating the mean abundance of each species across all longitudinal timepoints.

Beta diversity was evaluated by computing pairwise Aitchison distances between individuals. Principal Coordinate Analysis (PCoA) was performed on the resulting distance matrix to visualize the average community structure across conditions (non-IBD, CD, and UC) and clinical metadata, including sex, age category, and antibiotic usage (categorized by the percentage of timepoints with reported antibiotic exposure). Sex and antibiotic information were unavailable for the validation dataset.

Alpha diversity was assessed using the Shannon diversity index, calculated for each longitudinal sample and averaged across timepoints for each individual. Individual-level mean Shannon diversity was compared across disease conditions and, in the discovery dataset, further stratified by age category, sex, and antibiotic usage categories.

### Construction of individual-level microbial association networks

To construct individual-level MANs, we first applied a prevalence filter to each dataset, retaining species with an estimated read count greater than zero in at least 5% of samples. To minimize spurious network inference, individuals with fewer than seven longitudinal samples were excluded. After filtering, the discovery dataset comprised 165 species in 106 individuals, and the validation dataset comprised 224 species in 25 individuals.

Next, the abundance table was partitioned by individual, and species with zero total counts within a given individual were removed. We applied SparCC (Friedman and Alm, 2012) to each individual’s table, using a customized implementation based on the sparcc function of the R package SpiecEasi (v1.1.1) (Kurtz et al., 2015). To mitigate temporal dependencies between samples, we modified SparCC to detrend the data in CLR space before computing the log-ratio variances. Specifically, the CLR-transformed abundance of each species was regressed linearly on the position of the sample in the time-ordered series, and the residuals were used to compute the covariance matrix from which the variance of pairwise log-ratios was derived. SparCC was run with 100 outer and 30 inner iterations.

The resulting fully connected, undirected networks, were sparsified by removing edges with absolute SparCC weight below 0.3, 0.4, or 0.5. Finally, nodes whose species-level taxonomic label was marked as unclassified by the Pasolli et al. (2017) pipeline were removed after sparsification, yielding the final individual-level MANs.

### Node attributes

#### Taxonomic attributes

For each node (species), a binary taxonomic vector was initialized with positions corresponding to all unique taxa across the phylum, class, order, family, and genus levels observed in the discovery dataset. This vector was populated with a value of 1 at positions representing the specific taxa to which the species belonged, and 0 otherwise. For species exclusive to the validation dataset, the vector preserved any higher-level taxonomic ranks shared with the discovery dataset. For example, if *Bacteroides nordii* was absent from the discovery dataset, its vector in the validation set still incorporated its genus (*Bacteroides*) and all higher-level taxa that were present during training on the discovery dataset, setting unobserved taxa to 0.

#### Abundance attributes

To obtain individual-specific abundance attributes, we calculated the mean CLR-transformed abundance and the natural log-transformed temporal variance across longitudinal samples for each species within an individual. A pseudocount of 1 was added to estimated counts prior to CLR transformation to prevent undefined values. Additionally, a small numerical constant (*ɛ* = 10^−9^) was added prior to log-transforming variance to stabilize species with zero temporal variation. The resulting mean and variance values were concatenated into a bivariate vector capturing each species’ average abundance and stability.

#### Dummy attributes

Dummy attributes were created by assigning a univariate vector with a fixed value of 1 to all nodes across all individual networks in both the discovery and validation datasets. These attributes were used to evaluate model performance when relying strictly on network structure, isolated from any informative node attribute.

### Network metrics

Throughout the study, we quantified the structural properties of the individual-level MANs by computing several network metrics capturing node-, edge-, and global-level topological features:

#### Node degree

The number of edges incident to a node.

#### Normalized node degree

The node degree divided by *N −* 1, where *N* is the number of nodes in the MAN. This scales degree between 0 and 1, making it comparable across MANs with different numbers of nodes.

#### Node betweenness centrality

The number of shortest paths between all other pairs of nodes that pass through a given node.

#### Normalized node betweenness centrality

The node betweenness divided by (*N −* 1)(*N −* 2)*/*2, the number of node pairs excluding the node, scaling values between 0 and 1 for comparison across MANs of different sizes.

#### Number of nodes

The total number of bacterial species nodes present in the network.

#### Number of edges

The total number of associations retained in the network.

#### Positive-to-negative ratio

The ratio of the number of positive associations to the number of negative associations.

#### Density

The ratio of the actual number of edges to the maximum possible number of edges.

#### Assortativity

The Pearson correlation coefficient of degree between connected pairs of nodes, quantifying the tendency of nodes to connect with others of similar degree.

#### Global efficiency

A measure of how efficiently nodes are connected through short paths, based on the inverse of shortest path distances between node pairs. Edge weights (*w*) were converted to distances (*d*) using *d_ij_* = <u>^1^</u> , ensuring that stronger associations corresponded to shorter topological distances. Global efficiency was calculated as:

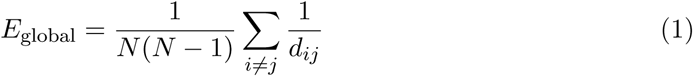

where *N* is the total number of nodes and *d_ij_*is the shortest path distance between nodes *i* and *j*.

#### P90/P50 degree ratio

The ratio between the 90^th^ percentile and the median (50^th^ percentile) of the node degree distribution. This metric quantifies hub dominance relative to median connectivity across the network.

Additionally, we assessed the similarity between individual-level MANs across taxonomic resolutions by calculating pairwise Jaccard similarity indices for both node and edge content. Nodes were progressively re-labeled from species to broader taxonomic categories (genus, family, order, class, phylum, and kingdom). For every pair of individuals within a given clinical condition (non-IBD or IBD), Jaccard similarities were computed between node sets (unique taxa present) and edge sets (unique associations).

### Training of traditional machine learning models

To establish a baseline for comparison with graph neural networks (GNNs), traditional machine learning models were trained on tabular taxonomic abundances at species level. Input counts were first normalized using CLR transformation. To make comparisons fair and aligned with individual-level MANs, longitudinal samples were aggregated to individuals by computing the mean of CLR-transformed abundances.

Random Forest (RF), Logistic Regression (LR), K-Nearest Neighbors (KNN), Support Vector Machine (SVM), and Multi-Layer Perceptron (MLP) models were trained for binary classification of non-IBD and IBD individuals. Each model was trained 50 times under a Monte Carlo-like approach where random 80:20 (train:test) dataset splits without replacement were used for each training-test run. The same random split seeds were used across all models for fair comparison. When applicable, class weights inversely proportional to class frequencies were applied to address class imbalance.

For each model and run, the training set was used for hyperparameter tuning via 3-fold crossvalidation. During tuning, data was Z-scaled and features with zero variance across individuals were removed. Important features were retained in the dataset based on ranking by ANOVA F-statistic values. The number of retained features could vary among 20%, 50%, 80%, or 100% of the original feature set. Hyperparameter grids for specific model types can be found in Supplementary Table 1. The best hyperparameter set for each model and run was selected by maximizing the area under the precision-recall curve (AUPRC). The model with the selected hyperparameters was then refit on the full training set and evaluated on the held-out test set (internal validation) and on all individuals of the validation dataset (external validation).

### Feature importance extraction from Random Forest models

Random Forest feature importance scores were extracted from the 50 Monte Carlo-like runs, retaining only runs with a held-out AUPRC *≥* 0.5 (n = 33). Within each run, importance scores were max-normalized, and species excluded by that run’s feature selection step were assigned an importance of zero. The final importance of each species was computed as its mean normalized importance across the retained runs, and the 20 highest-ranked species were selected for comparison with the prioritized nodes from GNNs.

### Training of graph neural networks

GNNs were trained using an architecture composed of two parallel, identical processing streams, following the separation of positive and negative relations commonly adopted in signed graph representation learning (Zhang et al., 2024). For each individual-level MAN, the network was divided into two subnetworks: a positive subnetwork comprising positive edges only, and a negative subnetwork comprising negative edges with their weights converted to absolute values. Before processing, edge weights in each subnetwork were rescaled to the [0, 1] interval by min-max normalization, with the minimum and maximum values estimated on the training graphs of each split and applied unchanged to the held-out and validation-dataset graphs to prevent data leakage, ensuring numerical stability during training. Self-loops were added to all nodes in both subnetworks so that isolated nodes could contribute to training. In the positive subnetwork, self-loops were assigned a weight of 1.0, whereas in the negative subnetwork they were assigned a weight of 0, reflecting the premise that a species cannot be negatively associated with itself. This yielded the weighted adjacency matrices **Ã**^+^ and **Ã**^−^, which shared the same node attribute matrix **X**.

Each subnetwork was processed by an independent stream, Φ^+^ or Φ^−^, consisting of a GraphConv (Morris et al., 2019) layer with additive aggregation, followed by GraphNorm (Cai et al., 2021) normalization, a ReLU activation function, and a dropout layer (rate = 0.3), and a second GraphConv layer. All convolutional layers had an output dimension of 128. The resulting node representations were then coarsened by a graph pooling layer (DiffPool) (Ying et al., 2018), in which two dedicated GraphConv layers computed the cluster assignment matrix with a compression ratio of 0.7, followed by global mean pooling to obtain a graph-level embedding per stream. All layers used their PyTorch Geometric implementations (Fey and Lenssen, 2019). The embeddings from both streams were concatenated and passed to a final linear layer to yield the logit:

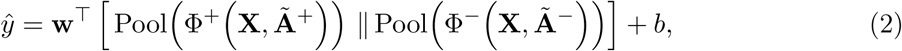

where Pool denotes DiffPool followed by global mean pooling, ‖ denotes concatenation, and **w** and *b* are the parameters of the linear layer. Because the two streams share no parameters or messages before concatenation, positive and negative associations are encoded independently and cannot cancel each other out during message passing.

GNNs were trained using a strategy similar to that of the traditional models, employing a Monte Carlo-like approach with random 80:20 train-test splits without replacement on the discovery dataset, using the same random split seeds across all models for a fair comparison. Models were trained across 50 independent iterations for binary classification (non-IBD vs. IBD) using a binary cross-entropy (BCE) loss with a class weight of 3.0 to address class imbalance, and optimized with the Adam optimizer at a learning rate of 5 *×* 10^−6^ without weight decay. Training ran for a maximum of 500 epochs, with early stopping based on internal validation loss and a patience of 50 epochs. Variations in node attributes, sparsification thresholds, and other parameters are summarized in Supplementary Table 2, and fixed configurations are listed in Supplementary Table 3.

### Interpretation of graph neural networks via Integrated Gradients

To identify network-level features relevant for classification, we trained 50 GNN model instances on the discovery dataset, each on a different stratified 80:20 train-test split. All instances used the dual-stream architecture with taxonomic and abundance node attributes, and MANs sparsified at an association weight threshold of 0.3. Training followed the procedure described above, except that dropout was set to 0 to stabilize attributions. To allow attribution to edge weights, the interpretation models received the signed SparCC weights as an input to the forward pass and split them into the positive and negative streams by rectification, assigning zero weight to opposite-sign edges in each stream. As a result, the DiffPool assignment layers, which do not use edge weights, operated on the full MAN topology in both streams.

We applied the Integrated Gradients algorithm (Sundararajan et al., 2017), as implemented in Captum (Kokhlikyan et al., 2020), to every MAN in the discovery dataset, including those used for training. Integrated Gradients estimates the importance of input elements by integrating the gradients of the model output with respect to the input along a linear path from a baseline to the original input, approximated in *m* steps:

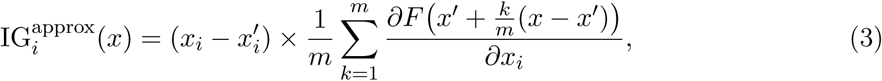

where *x_i_* is the original value of input element *i*, *x*^′^*_i_* its baseline value, and *F* the model output logit. Attributions were computed with *m* = 200 steps. Because non-IBD individuals were encoded as the positive class, positive attributions indicate contributions toward the non-IBD class and negative attributions toward the IBD class.

Node and edge attributions were computed in separate passes. For node attributes, edge weights were held at their original values, and baselines were set to represent the absence or minimal state of biological information: zero vectors for taxonomic attributes, and the 5th percentile of all observed values for the mean CLR-transformed abundance and its variance, since zero does not denote absence in CLR-transformed data. For edges, attributions were computed with respect to the signed SparCC weights, with node attributes held fixed and a baseline weight of zero, corresponding to the absence of an association.

For each model instance, predicted probabilities for all discovery MANs were used to compute a pooled AUPRC, and only instances with a pooled AUPRC *≥* 0.5 were retained (n = 30) for downstream analyses. Because these values include training samples, the threshold served only to exclude poorly fitting models and not to estimate generalization performance.

### Node importance

For each retained model instance *r*, MAN *g*, and node *i*, attributions were summed across all attributes *j* of the node to obtain a net (signed) and an absolute importance score:

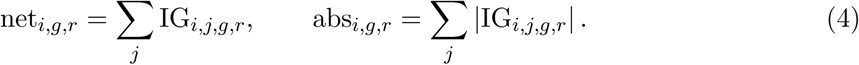

Both scores were averaged across instances, and the directional stability of each node was computed as *|*Σ*_r_* net*_i,g,r_| /* Σ*_r_ |*net*_i,g,r_|*, ranging from 0 (signs inconsistent across instances) to 1 (signs fully consistent). Averaged scores were then normalized within each MAN by dividing by the MAN’s maximum absolute node importance. At the cohort level, the weighted node importance (WNI*_i_*) was defined as the mean normalized absolute importance across the MANs containing node *i*, multiplied by its prevalence (the proportion of MANs containing the node), and the 20 nodes with the highest WNI*_i_* were prioritized. To assess the contribution of each attribute type, absolute attributions were also summed separately over taxonomic and abundance attributes, averaged and normalized as above, and expressed as a proportion of the node’s absolute importance.

### Edge importance and importance metagraph

Because each undirected association is represented by two directed edges in the model input (Fey and Lenssen, 2019), the attributions of both directions were summed to obtain a single signed importance per association and model instance, and its absolute value was taken as the absolute importance. Signed and absolute edge scores were averaged across instances, normalized within each MAN by the MAN’s maximum absolute edge importance, and summarized as a weighted edge importance (WEI*_e_*), defined analogously to WNI*_i_*. The importance metagraph was defined as the largest connected component of the network formed by the 100 edges with the highest WEI*_e_* and their incident nodes. Taxonomic cohesion of the metagraph was quantified, at each rank from kingdom to genus, as the proportion of metagraph nodes belonging to the most common taxon.

### Ablation experiments

To assess the contribution of the metagraph to classification, we constructed two variants of every individual-level MAN in both datasets: metagraph-only graphs (M only), reduced to the module induced by metagraph nodes, and removal graphs (WG - M), in which metagraph nodes and their incident edges were removed from the whole graph (WG). In the validation dataset, metagraph nodes were identified by matching species names to the metagraph defined in the discovery dataset. To separate the contribution of associations from that of node attributes, we also generated an edge-free version of each configuration, in which all edges were removed while nodes and node attributes were kept.

For each of the six configurations, models were trained on the discovery dataset as described above (taxonomic and abundance attributes; sparsification threshold of 0.3) over 50 Monte Carlo-like runs with stratified 80:20 splits shared across configurations. Each trained model was evaluated on the validation dataset in all six configurations, and performance was summarized as the mean and standard deviation of AUPRC and AUROC across runs.

### Analysis of Bacteroidales SparCC weights

To characterize the associations underlying the predictive signal of the metagraph, we analyzed the SparCC weights among the 17 metagraph species belonging to the order Bacteroidales, using the sparsified MANs of both datasets (absolute association weight threshold of 0.3). For each individual, we enumerated all pairs of these species present in their MAN and assigned each pair the weight of the corresponding edge, or zero when the two species were not connected.

At the individual level, we computed the mean weight across all pairs and compared non-IBD and IBD individuals within each dataset. At the pair level, we computed the mean weight separately in non-IBD and IBD individuals and their difference, restricting the comparison to pairs present in at least three individuals of each condition.

### Statistical analyses

Statistical comparisons were performed using non-parametric tests, which do not require assumptions about the distribution of the data. Differences between groups (e.g., non-IBD vs. IBD individuals) in diversity, abundance, association weights, and network metrics were assessed with two-sided Mann-Whitney U tests, except for Shannon diversity between non-IBD and IBD individuals, where a one-sided test was used to evaluate the expected reduction of diversity in IBD. Differences in node prevalence across conditions were assessed with two-sided Fisher’s exact tests. Performance differences between ablation configurations were assessed with two-sided Wilcoxon signed-rank tests paired by model run, since all configurations shared the same train-test splits. Relationships between node importance, centrality, and abundance were evaluated with Spearman’s rank correlation. Whenever multiple tests were performed, *p*-values were adjusted with the Benjamini–Hochberg procedure. Results were considered significant at *p <* 0.05 for single comparisons and at FDR *<* 0.05 for corrected comparisons.

## Data and code availability

The metagenomic datasets analyzed in this study are publicly available through the curatedMetagenomicData R package (v3.14.0) under the study names “HMP_2019_ibdmdb” (Lloyd-Price et al., 2019) and “HallAB_2017” (Hall et al., 2017). All code used for data processing, network inference, model training, interpretation, and figure generation is available on GitHub at https://github.com/Multiomics-Analytics-Group/IBD_Individual_MANs. The raw and filtered abundance tables, individual-level microbial association networks, node attribute tables, PyTorch Geometric datasets, model performance results, Integrated Gradients attributions, and ablation results are deposited in Zenodo (https://doi.org/10.5281/zenodo.23213629).

## Acknowledgments

This work was supported by the Novo Nordisk Foundation (NNF25SA0109652; NNF24SA0100980; NNF21OC0069089) and Calmette & Yersin PhD Grant (Pasteur Network, Sep/2023)

## Declaration of generative AI in the writing process

During the preparation of the manuscript, the authors used Claude (Anthropic) to assist with language polishing. After using this tool, the authors reviewed and edited the text as needed and take full responsibility for the content of this work.

## Supplementary figures

**Figure S1:**
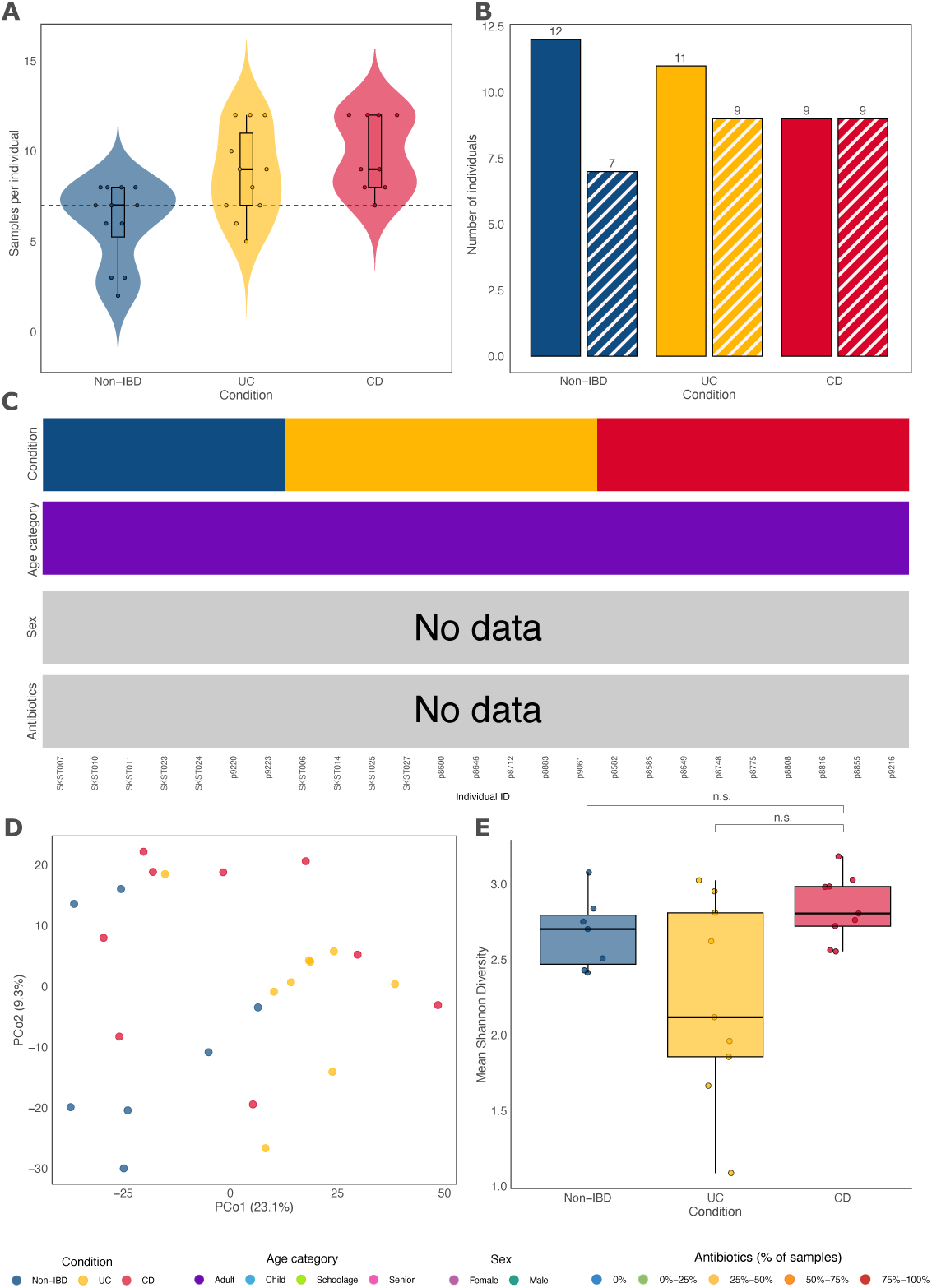
Overview and microbial diversity analysis of the validation dataset. (**A**) Distribution of the number of longitudinal samples per individual. The dashed line indicates the threshold (*<* 7 samples) used to filter out individuals. (**B**) Number of individuals in the validation dataset before (solid fill) and after (hatched fill) filtering using the threshold shown in A. (**C**) Visual representation of individual metadata, including condition and age category. Sex and antibiotic usage information is unavailable for the validation dataset. (**D**) Principal Coordinate Analysis (PCoA) ordination of Aitchison distances between individuals computed on mean CLR-transformed species abundances. (**E**) Box plots showing the distribution of mean Shannon diversity values within each condition. Statistical significance was not reached either by comparing non-IBD and IBD (one-sided Mann-Whitney U test, *p* = 0.559) or UC and CD (two-sided Mann-Whitney U test, *p* = 0.062). n.s., non-significant.

**Figure S2:**
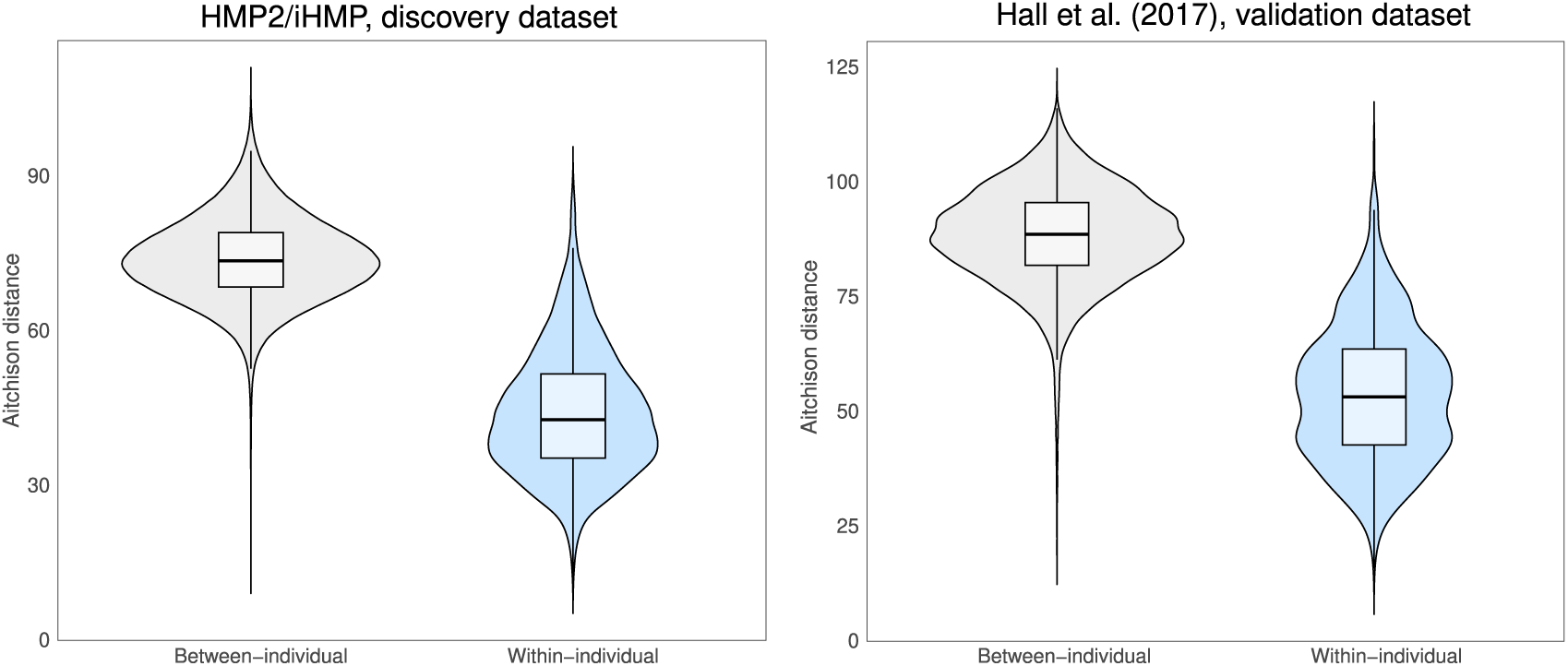
Evaluation of between- and within-individual distances in the discovery and validation datasets. Pairwise Aitchison distances were computed across longitudinal samples to evaluate the overall stability of individual microbiota compositions relative to inter-individual variation.

**Figure S3:**
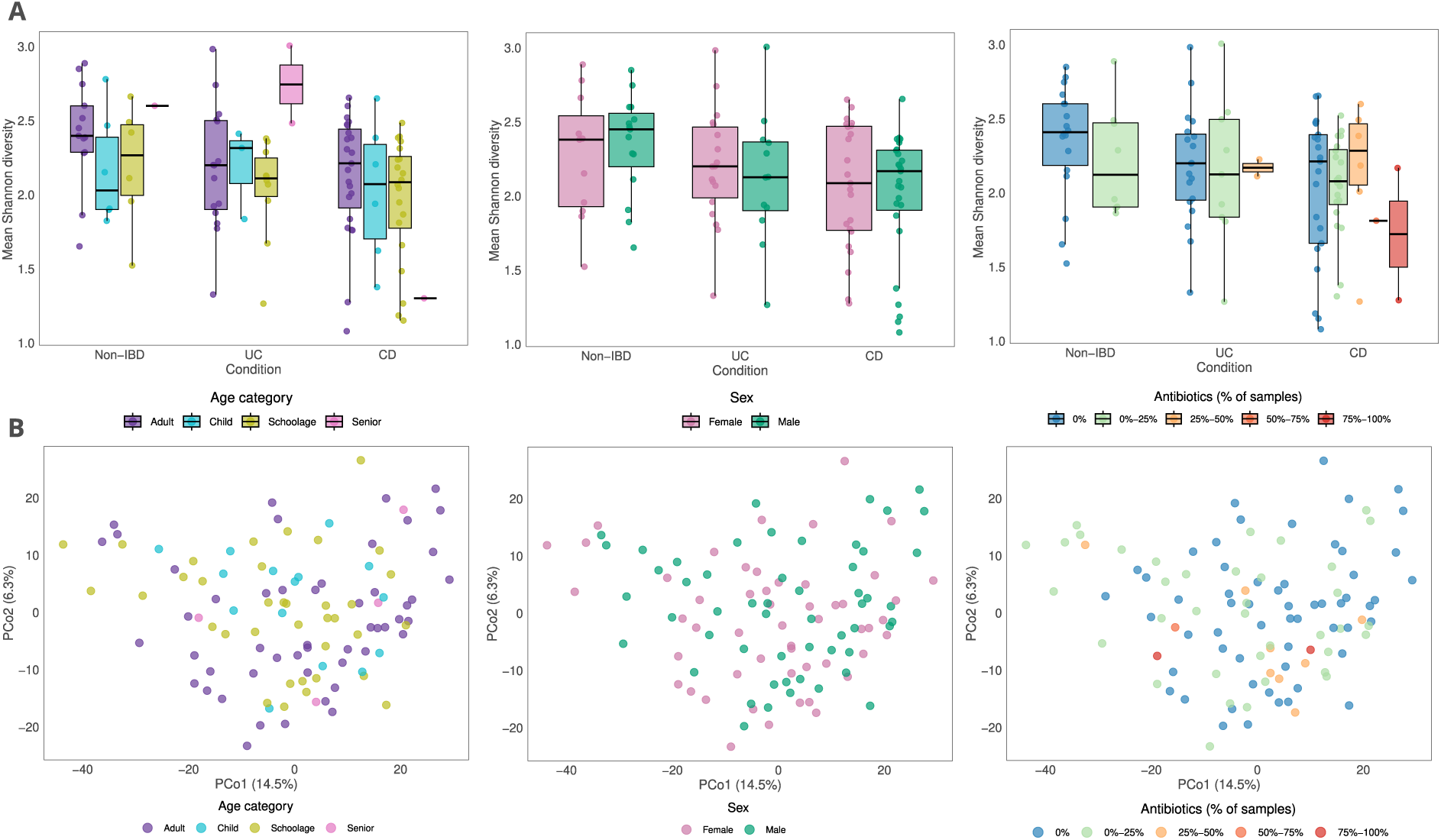
Stratified microbial diversity analysis of the discovery dataset. (**A**) Box plots showing the distribution of mean Shannon diversity values stratified by age group, sex, and the percentage of timepoints where antibiotic usage was reported. (**B**) Principal Coordinate Analysis (PCoA) ordinations based on Aitchison distances, colored by age, sex, and antibiotic usage. Aitchison distances were computed individual-wise using mean CLR-transformed species abundances.

**Figure S4:**
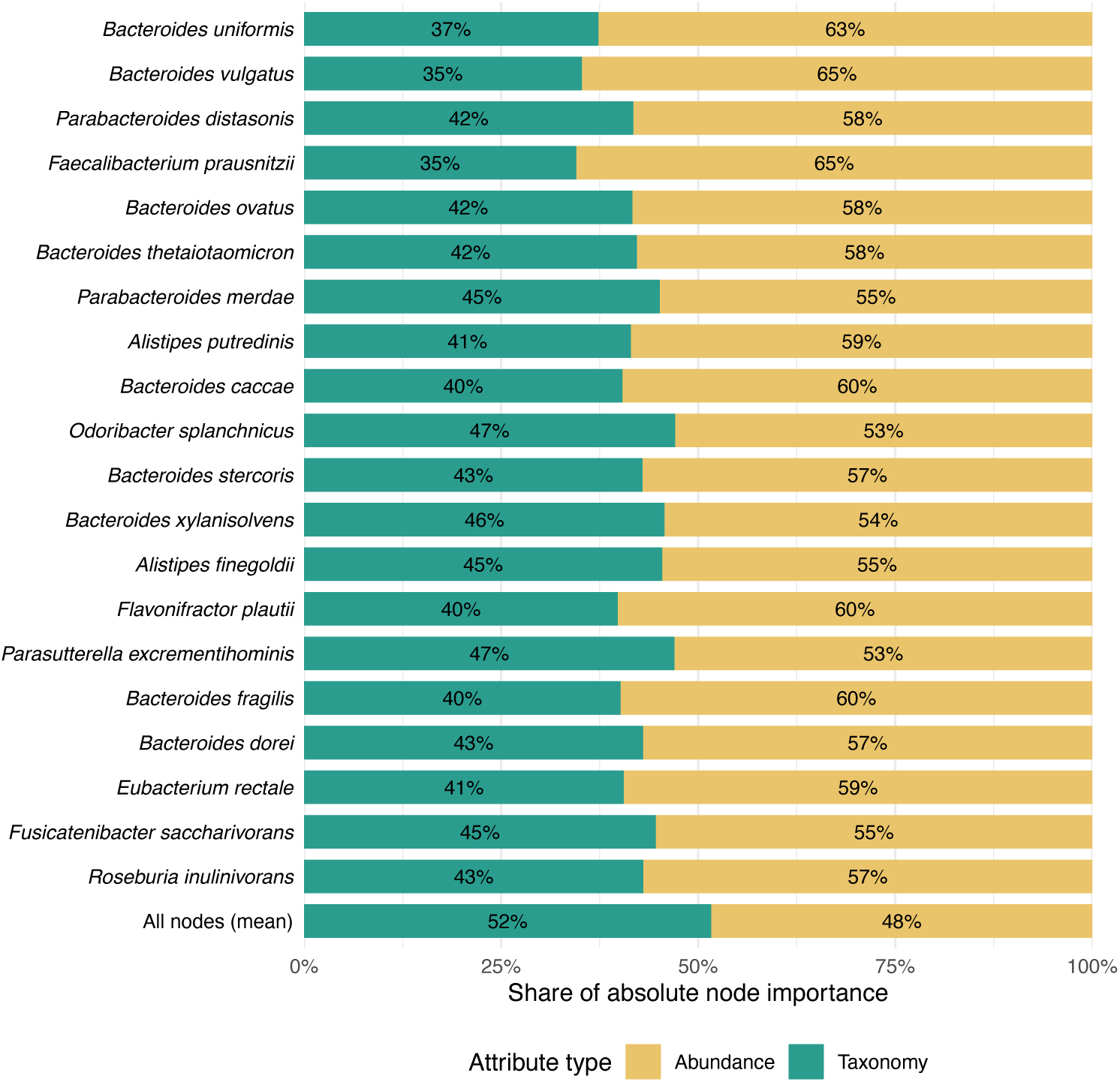
Contribution of node attribute types to absolute node importance. Share of absolute node importance assigned to taxonomic and abundance attributes (mean CLR-transformed abundance and log-transformed variance) for the top 20 prioritized nodes, ranked by prevalence-weighted mean absolute importance (as in Fig. 4), and the mean across all nodes in the discovery dataset (bottom bar). For each node, absolute Integrated Gradients scores were summed within each attribute type, averaged across model instances and MANs, and expressed as a proportion of the node’s absolute importance.

**Figure S5:**
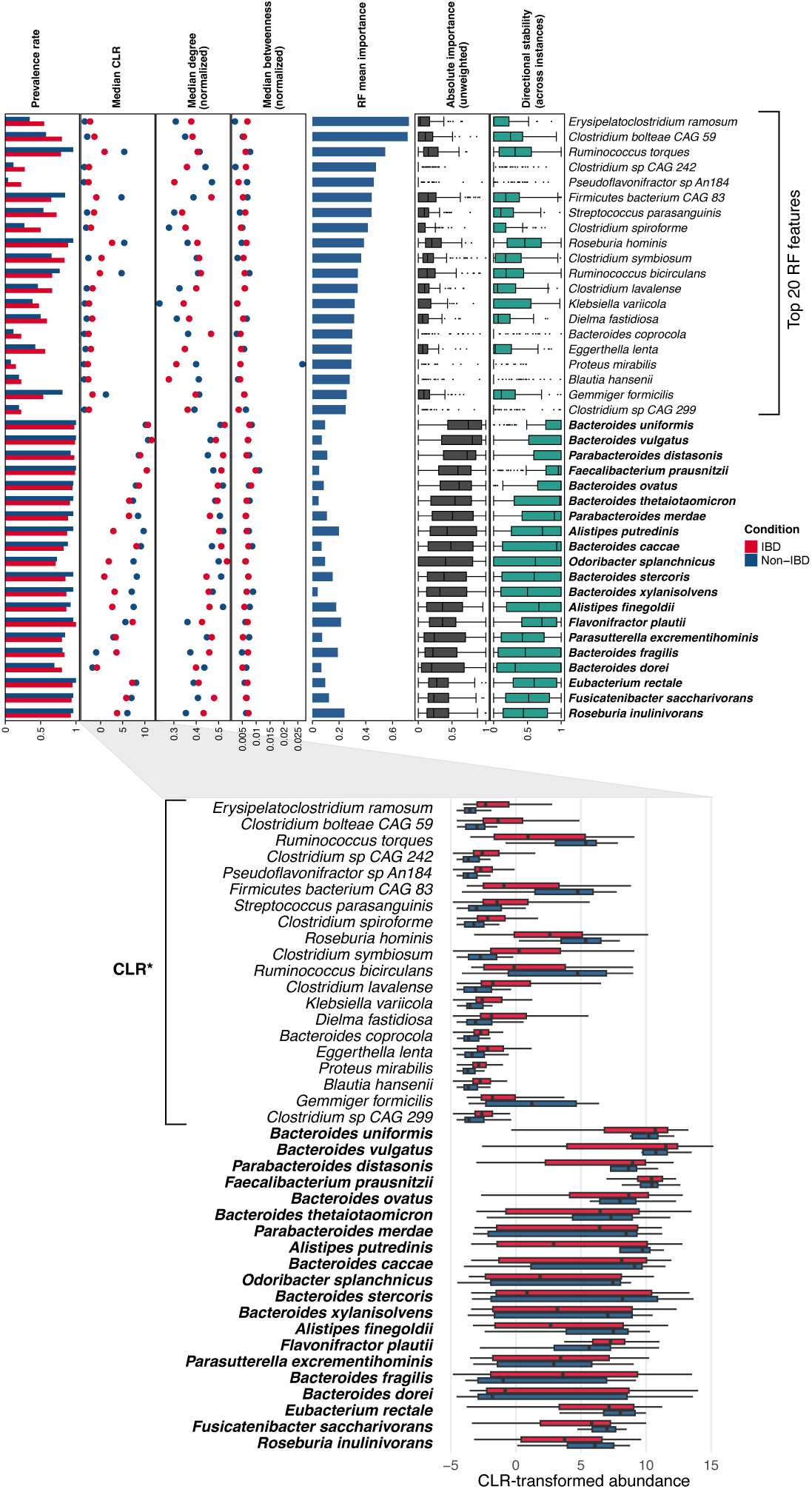
Comparison of top-ranked features from Random Forest models and prioritized nodes from GNNs. Top-ranked Random Forest (RF) features were extracted from 33 well-fitting model instances (AUPRC *≥* 0.50, computed on held-out only) across 50 Monte Carlo-like iterations. RF importance scores were max-normalized within each model instance, and the mean was calculated for each species across all instances to yield the final RF mean importance score. Top panel: The top 20 RF features are ranked by this score, followed by top prioritized nodes (indicated in bold) ranked by prevalence-weighted mean absolute importance scores (as in Fig. 4). Left annotations display the prevalence rate, median CLR-transformed abundance, median normalized degree, and betweenness per condition. Right annotations display the distribution of node absolute importance and signed importance directional stability. Node metrics were computed for both top prioritized nodes and RF features (represented as nodes in individual-level MANs) for comparison. Bottom panel: Distributions of CLR-transformed abundances for top-ranked RF features and prioritized nodes from GNNs. Asterisks indicate statistical significance from two-sided Mann-Whitney U tests after Benjamini–Hochberg FDR correction (FDR < 0.05)

**Figure S6:**
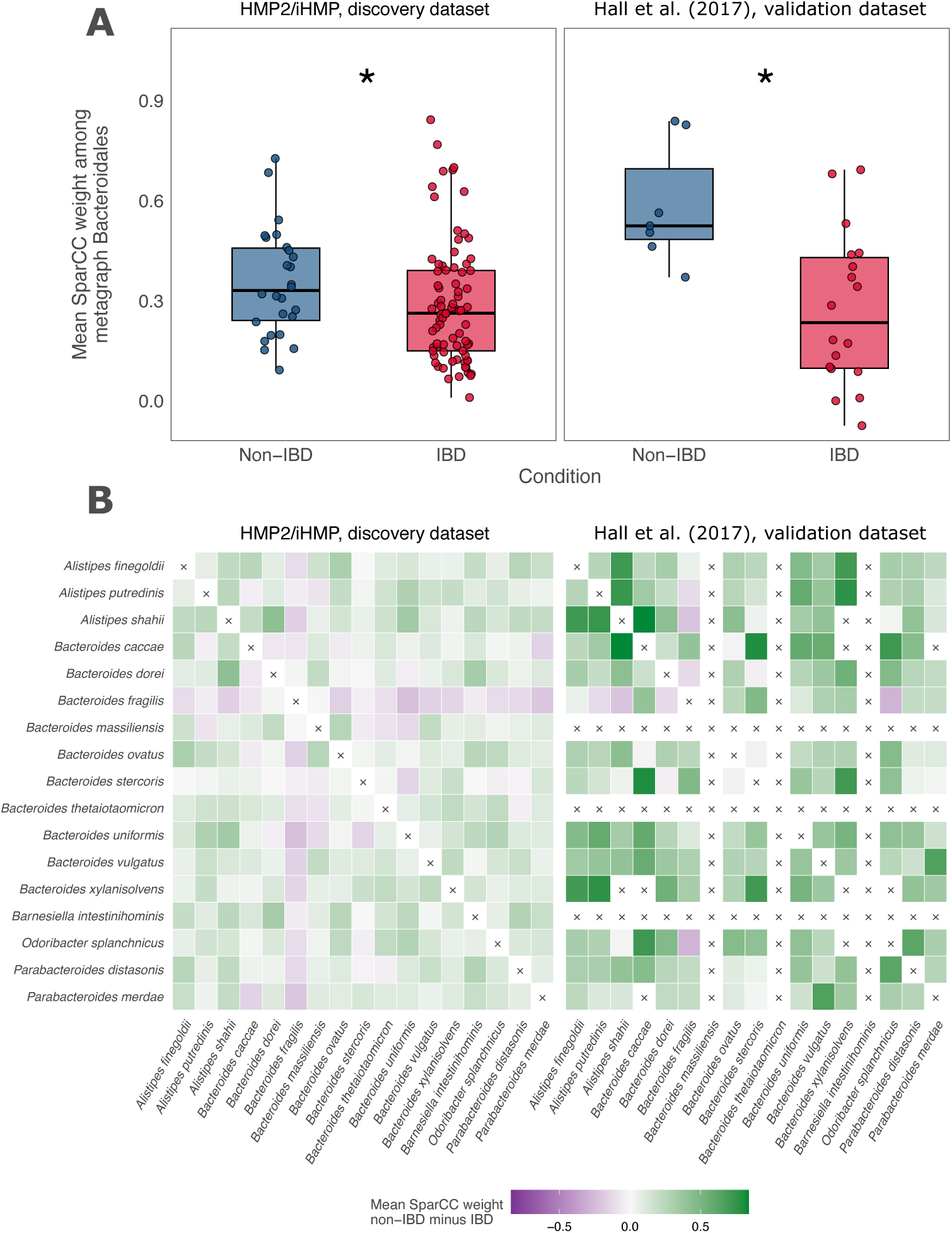
Associations among metagraph Bacteroidales in non-IBD and IBD individual-level MANs. (**A**) Mean SparCC weight among the 17 Bacteroidales species of the metagraph, computed for each individual across all pairs of these species present in their MAN, with pairs not connected by an edge counted as zero. Each point represents one individual (*, Mann-Whitney U test, *p* < 0.05). (**B**) Difference in mean SparCC weight between non-IBD and IBD individuals (non-IBD minus IBD) for each pair of metagraph Bacteroidales, in the discovery (left) and validation (right) datasets (green, higher in non-IBD, and purple, higher in IBD). Associations were taken from the sparsified MANs used as model input (absolute association weight threshold of 0.3), with pairs not connected by an edge counted as zero. Pairs of species co-occurring in fewer than three individuals of either condition were not evaluated. These pairs and the diagonal are marked with ×.

## References

1. F. Backhed, R. E. Ley, J. L. Sonnenburg, D. A. Peterson, and J. I. Gordon. Host-bacterial mutualism in the human intestine. Science, 307(5717):1915–1920, 2005.

2. B. Bakir-Gungor, H. Hacılar, A. Jabeer, O. U. Nalbantoglu, O. Aran, and M. Yousef. Inflammatory bowel disease biomarkers of human gut microbiota selected via different feature selection methods. PeerJ, 10:e13205, 2022.

3. F. Beghini, L. J. McIver, A. Blanco-Míguez, L. Dubois, F. Asnicar, S. Maharjan, A. Mailyan, P. Manghi, M. Scholz, A. M. Thomas, et al. Integrating taxonomic, functional, and strain-level profiling of diverse microbial communities with biobakery 3. elife, 10:e65088, 2021.

4. G. Berg, D. Rybakova, D. Fischer, T. Cernava, M.-C. C. Vergès, T. Charles, X. Chen, L. Cocolin, K. Eversole, G. H. Corral, et al. Microbiome definition re-visited: old concepts and new challenges. Microbiome, 8(1):103, 2020.

6. T. C. Bosch and M. J. McFall-Ngai. Metaorganisms as the new frontier. Zoology, 114(4):185–190, 2011.

7. R. M. Braga and R. L. Buckner. Parallel interdigitated distributed networks within the individual estimated by intrinsic functional connectivity. Neuron, 95(2):457–471, 2017.

8. T. Cai, S. Luo, K. Xu, D. He, T.-y. Liu, and L. Wang. Graphnorm: A principled approach to accelerating graph neural network training. In International Conference on Machine Learning, pages 1204–1215. PMLR, 2021.

9. R. Caruso, B. C. Lo, and G. Núñez. Host–microbiota interactions in inflammatory bowel disease. Nature reviews immunology, 20(7):411–426, 2020.

10. H.-H. Chen, C.-W. Hsueh, C.-H. Lee, T.-Y. Hao, T.-Y. Tu, L.-Y. Chang, J.-C. Lee, and C.-Y. Lin. Sweet: a single-sample network inference method for deciphering individual features in disease. Briefings in bioinformatics, 24(2):bbad032, 2023.

11. L. Chen, V. Collij, M. Jaeger, I. C. van den Munckhof, A. Vich Vila, A. Kurilshikov, R. Gacesa, T. Sinha, M. Oosting, L. A. Joosten, et al. Gut microbial co-abundance networks show specificity in inflammatory bowel disease and obesity. Nature communications, 11(1):4018, 2020.

12. J. Chow, H. Tang, and S. K. Mazmanian. Pathobionts of the gastrointestinal microbiota and inflammatory disease. Current opinion in immunology, 23(4):473–480, 2011.

13. M. De Marzio, K. Glass, and M. L. Kuijjer. Single-sample network modeling on omics data. BMC biology, 21(1):296, 2023.

14. K. Faust and J. Raes. Microbial interactions: from networks to models. Nature reviews microbiology, 10(8):538–550, 2012.

15. K. Faust, J. F. Sathirapongsasuti, J. Izard, N. Segata, D. Gevers, J. Raes, and C. Huttenhower. Microbial co-occurrence relationships in the human microbiome. PLoS computational biology, 8(7):e1002606, 2012.

16. M. Fey and J. E. Lenssen. Fast graph representation learning with pytorch geometric. *arXiv preprint arXiv:1903.02428*, 2019.

17. E. S. Finn, X. Shen, D. Scheinost, M. D. Rosenberg, J. Huang, M. M. Chun, X. Papademetris, and R. T. Constable. Functional connectome fingerprinting: identifying individuals using patterns of brain connectivity. Nature neuroscience, 18(11):1664–1671, 2015.

18. J. Friedman and E. J. Alm. Inferring correlation networks from genomic survey data. PLoS computational biology, 8(9), 2012.

19. T. Geese, C. Bang, A. Franke, W. Lieb, and A. Dempfle. The human gut microbiota in ibd, characterizing hubs, the core microbiota and terminal nodes: a network-based approach. BMC microbiology, 25(1):371, 2025.

20. D. Gevers, S. Kugathasan, L. A. Denson, Y. Vázquez-Baeza, W. Van Treuren, B. Ren, E. Schwager, D. Knights, S. J. Song, M. Yassour, et al. The treatment-naive microbiome in new-onset crohn’s disease. Cell host & microbe, 15(3):382–392, 2014.

21. E. Z. Gomaa. Human gut microbiota/microbiome in health and diseases: a review. Antonie van Leeuwenhoek, 113(12):2019–2040, 2020.

22. C. Gratton, T. O. Laumann, A. N. Nielsen, D. J. Greene, E. M. Gordon, A. W. Gilmore, S. M. Nelson, R. S. Coalson, A. Z. Snyder, B. L. Schlaggar, et al. Functional brain networks are dominated by stable group and individual factors, not cognitive or daily variation. Neuron, 98 (2):439–452, 2018.

23. A. B. Hall, M. Yassour, J. Sauk, A. Garner, X. Jiang, T. Arthur, G. K. Lagoudas, T. Vatanen, N. Fornelos, R. Wilson, et al. A novel ruminococcus gnavus clade enriched in inflammatory bowel disease patients. Genome medicine, 9(1):103, 2017.

24. K. Honda and D. R. Littman. The microbiome in infectious disease and inflammation. Annual review of immunology, 30:759–795, 2012.

25. X. Hu, L. Chu, J. Pei, W. Liu, and J. Bian. Model complexity of deep learning: A survey. Knowledge and information systems, 63(10):2585–2619, 2021.

26. V. A. Ivanov, W. H. Hartman, and M. Soheilypour. Decoding the microbiome-disease axis with interpretable graph neural networks. Journal of Applied Microbiology, 137(4):lxag063, 2026.

27. S. Kodikara, S. Ellul, and K.-A. Lê Cao. Statistical challenges in longitudinal microbiome data analysis. Briefings in Bioinformatics, 23(4):bbac273, 2022.

28. N. Kokhlikyan, V. Miglani, M. Martin, E. Wang, B. Alsallakh, J. Reynolds, A. Melnikov, N. Kliushkina, C. Araya, S. Yan, et al. Captum: A unified and generic model interpretability library for pytorch. *arXiv preprint arXiv:2009.07896*, 2020.

29. M. Kuhn, T. S. B. Schmidt, P. Ferretti, A. Głazek, S. M. Robbani, W. Akanni, A. Fullam, C. Schudoma, E. Cetin, M. Hassan, et al. Metalog: curated and harmonised contextual data for global metagenomics samples. Nucleic acids research, 54(D1):D826–D834, 2026.

30. M. L. Kuijjer, M. G. Tung, G. Yuan, J. Quackenbush, and K. Glass. Estimating sample-specific regulatory networks. Iscience, 14:226–240, 2019.

31. Z. D. Kurtz, C. L. Müller, E. R. Miraldi, D. R. Littman, M. J. Blaser, and R. A. Bonneau. Sparse and compositionally robust inference of microbial ecological networks. PLoS computational biology, 11(5):e1004226, 2015.

32. T. J. Lam and Y. Ye. Meta-analysis of microbiome association networks reveal patterns of dysbiosis in diseased microbiomes. Scientific Reports, 12(1):17482, 2022.

33. A. Lamurias, M. Sereika, M. Albertsen, K. Hose, and T. D. Nielsen. Metagenomic binning with assembly graph embeddings. Bioinformatics, 38(19):4481–4487, 2022.

34. Y. Lee, M. Cappellato, and B. Di Camillo. Machine learning–based feature selection to search stable microbial biomarkers: application to inflammatory bowel disease. GigaScience, 12: giad083, 2023.

35. Y. Lee, J. Seo, and B. Di Camillo. Shap-based binarization enhances metataxonomic machine learning with application to gut microbiota of inflammatory bowel disease. Scientific reports, 15(1):39059, 2025.

36. R. Leinonen, H. Sugawara, M. Shumway, and I. N. S. D. Collaboration. The sequence read archive. Nucleic acids research, 39(suppl_1):D19–D21, 2010.

37. P. Li, M. Li, and W.-H. Chen. Best practices for developing microbiome-based disease diagnostic classifiers through machine learning. Gut Microbes, 17(1):2489074, 2025.

38. J. Liñares-Blanco, C. Fernandez-Lozano, J. A. Seoane, and G. López-Campos. Machine learning based microbiome signature to predict inflammatory bowel disease subtypes. Frontiers in microbiology, 13:872671, 2022.

39. H. Liu, C. Liao, L. Wu, J. Tang, J. Chen, C. Lei, L. Zheng, C. Zhang, Y.-Y. Liu, J. Xavier, et al. Ecological dynamics of the gut microbiome in response to dietary fiber. The ISME journal, 16 (8):2040–2055, 2022.

40. I. Liu, Y. Wang, H. Ji, K. Aihara, and L. Chen. Personalized characterization of diseases using sample-specific networks. Nucleic acids research, 44(22):e164–e164, 2016.

41. J. Lloyd-Price, C. Arze, A. N. Ananthakrishnan, M. Schirmer, J. Avila-Pacheco, T. W. Poon, E. Andrews, N. J. Ajami, K. S. Bonham, C. J. Brislawn, et al. Multi-omics of the gut microbial ecosystem in inflammatory bowel diseases. Nature, 569(7758):655–662, 2019.

42. P. Louis and H. J. Flint. Diversity, metabolism and microbial ecology of butyrate-producing bacteria from the human large intestine. FEMS microbiology letters, 294(1):1–8, 2009.

43. C. Lupp, M. L. Robertson, M. E. Wickham, I. Sekirov, O. L. Champion, E. C. Gaynor, and B. B. Finlay. Host-mediated inflammation disrupts the intestinal microbiota and promotes the overgrowth of enterobacteriaceae. Cell host & microbe, 2(2):119–129, 2007.

44. A. V. Mardanov, V. V. Kadnikov, and N. V. Ravin. Metagenomics: A paradigm shift in microbiology. In Metagenomics, pages 1–15. Elsevier, 2025.

45. M. S. Matchado, M. Lauber, S. Reitmeier, T. Kacprowski, J. Baumbach, D. Haller, and M. List. Network analysis methods for studying microbial communities: A mini review. Computational and structural biotechnology journal, 19:2687–2698, 2021.

46. R. H. Mills, P. S. Dulai, Y. Vázquez-Baeza, C. Sauceda, N. Daniel, R. R. Gerner, L. E. Batachari, M. Malfavon, Q. Zhu, K. Weldon, et al. Multi-omics analyses of the ulcerative colitis gut microbiome link bacteroides vulgatus proteases with disease severity. Nature microbiology, 7 (2):262–276, 2022.

47. C. Morris, M. Ritzert, M. Fey, W. L. Hamilton, J. E. Lenssen, G. Rattan, and M. Grohe. Weisfeiler and leman go neural: Higher-order graph neural networks. In Proceedings of the AAAI conference on artificial intelligence, volume 33, pages 4602–4609, 2019.

48. A. Nishida, R. Inoue, O. Inatomi, S. Bamba, Y. Naito, and A. Andoh. Gut microbiota in the pathogenesis of inflammatory bowel disease. Clinical journal of gastroenterology, 11(1):1–10, 2018.

49. K. Nomura, D. Ishikawa, K. Okahara, S. Ito, K. Haga, M. Takahashi, A. Arakawa, T. Shibuya, T. Osada, K. Kuwahara-Arai, et al. Bacteroidetes species are correlated with disease activity in ulcerative colitis. Journal of clinical medicine, 10(8):1749, 2021.

50. A. Palleja, K. H. Mikkelsen, S. K. Forslund, A. Kashani, K. H. Allin, T. Nielsen, T. H. Hansen, S. Liang, Q. Feng, C. Zhang, et al. Recovery of gut microbiota of healthy adults following antibiotic exposure. Nature microbiology, 3(11):1255–1265, 2018.

51. S. Pan, X. Jiang, and K. Zhang. Wsgmb: weight signed graph neural network for microbial biomarker identification. Briefings in bioinformatics, 25(1):bbad448, 2024.

52. E. Pasolli, D. T. Truong, F. Malik, L. Waldron, and N. Segata. Machine learning meta-analysis of large metagenomic datasets: tools and biological insights. PLoS computational biology, 12 (7):e1004977, 2016.

53. E. Pasolli, L. Schiffer, P. Manghi, A. Renson, V. Obenchain, D. T. Truong, F. Beghini, F. Malik, M. Ramos, J. B. Dowd, et al. Accessible, curated metagenomic data through experimenthub. Nature methods, 14(11):1023–1024, 2017.

54. E. Quévrain, M. Maubert, C. Michon, F. Chain, R. Marquant, J. Tailhades, S. Miquel, L. Carlier, L. Bermúdez-Humarán, B. Pigneur, et al. Identification of an anti-inflammatory protein from faecalibacterium prausnitzii, a commensal bacterium deficient in crohn’s disease. Gut, 65(3): 415–425, 2016.

55. S. Rakoff-Nahoum, K. R. Foster, and L. E. Comstock. The evolution of cooperation within the gut microbiota. Nature, 533(7602):255–259, 2016.

56. G. P. Ramos and K. A. Papadakis. Mechanisms of disease: inflammatory bowel diseases. In Mayo Clinic Proceedings, volume 94, pages 155–165. Elsevier, 2019.

57. T. A. Rice, A. A. Bielecka, M. T. Nguyen, C. E. Rosen, D. Song, N. D. Sonnert, Y. Yang, Y. Cao, V. Khetrapal, J. R. Catanzaro, et al. Interspecies commensal interactions have nonlinear impacts on host immunity. Cell host & microbe, 30(7):988–1002, 2022.

58. L. Richardson, B. Allen, G. Baldi, M. Beracochea, M. L. Bileschi, T. Burdett, J. Burgin, J. Caballero-Pérez, G. Cochrane, L. J. Colwell, et al. Mgnify: the microbiome sequence data analysis resource in 2023. Nucleic acids research, 51(D1):D753–D759, 2023.

59. L. Rigottier-Gois. Dysbiosis in inflammatory bowel diseases: the oxygen hypothesis. The ISME journal, 7(7):1256–1261, 2013.

60. R. B. Sartor and G. D. Wu. Roles for intestinal bacteria, viruses, and fungi in pathogenesis of inflammatory bowel diseases and therapeutic approaches. Gastroenterology, 152(2):327–339, 2017.

61. O. Shtossel, H. Isakov, S. Turjeman, O. Koren, and Y. Louzoun. Ordering taxa in image convolution networks improves microbiome-based machine learning accuracy. Gut Microbes, 15 (1):2224474, 2023.

62. J.-C. Simon, J. R. Marchesi, C. Mougel, and M.-A. Selosse. Host-microbiota interactions: from holobiont theory to analysis. Microbiome, 7(1):5, 2019.

63. E. H. Simpson. The interpretation of interaction in contingency tables. Journal of the Royal Statistical Society: Series B (Methodological), 13(2):238–241, 1951.

64. H. Sokol, B. Pigneur, L. Watterlot, O. Lakhdari, L. G. Bermúdez-Humarán, J.-J. Gratadoux, S. Blugeon, C. Bridonneau, J.-P. Furet, G. Corthier, et al. Faecalibacterium prausnitzii is an anti-inflammatory commensal bacterium identified by gut microbiota analysis of crohn disease patients. Proceedings of the National Academy of Sciences, 105(43):16731–16736, 2008.

65. H. Sokol, P. Seksik, J. Furet, O. Firmesse, I. Nion-Larmurier, L. Beaugerie, J. Cosnes, G. Corthier, P. Marteau, and J. Doré. Low counts of faecalibacterium prausnitzii in colitis microbiota. Inflammatory bowel diseases, 15(8):1183–1189, 2009.

66. A. M. Spees, C. A. Lopez, D. D. Kingsbury, S. E. Winter, and A. J. Bäumler. Colonization resistance: battle of the bugs or ménage à trois with the host? PLoS pathogens, 9(11):e1003730, 2013.

67. I. S. Stafford, M. M. Gosink, E. Mossotto, S. Ennis, and M. Hauben. A systematic review of artificial intelligence and machine learning applications to inflammatory bowel disease, with practical guidelines for interpretation. Inflammatory Bowel Diseases, 28(10):1573–1583, 2022.

68. M. Sundararajan, A. Taly, and Q. Yan. Axiomatic attribution for deep networks. In International conference on machine learning, pages 3319–3328. PMLR, 2017.

69. K. Syama, J. A. A. Jothi, and N. Khanna. Automatic disease prediction from human gut metagenomic data using boosting graphsage. BMC bioinformatics, 24(1):126, 2023.

70. B. D. Topçuoğlu, N. A. Lesniak, M. T. Ruffin IV, J. Wiens, and P. D. Schloss. A framework for effective application of machine learning to microbiome-based classification problems. MBio, 11(3):10–1128, 2020.

71. M. Van Hul, P. D. Cani, C. Petitfils, W. M. De Vos, H. Tilg, and E. M. El-Omar. What defines a healthy gut microbiome? Gut, 73(11):1893–1908, 2024.

72. O. S. Venturelli, A. V. Carr, G. Fisher, R. H. Hsu, R. Lau, B. P. Bowen, S. Hromada, T. Northen, and A. P. Arkin. Deciphering microbial interactions in synthetic human gut microbiome communities. Molecular systems biology, 14(6):MSB178157, 2018.

73. G. Wei and Y. Liu. A graph contrastive learning method for enhancing genome recovery in complex microbial communities. Entropy, 27(9):921, 2025.

74. S. Weiss, W. Van Treuren, C. Lozupone, K. Faust, J. Friedman, Y. Deng, L. C. Xia, Z. Z. Xu, L. Ursell, E. J. Alm, et al. Correlation detection strategies in microbial data sets vary widely in sensitivity and precision. The ISME journal, 10(7):1669–1681, 2016.

75. Y. Xu, K. Nash, A. Acharjee, and G. V. Gkoutos. Caconet: a novel classification framework for microbial correlation networks. Bioinformatics, 38(6):1639–1647, 2022.

76. H. Xue, V. Mallawaarachchi, Y. Zhang, V. Rajan, and Y. Lin. Repbin: constraint-based graph representation learning for metagenomic binning. In *Proceedings of the AAAI Conference on Artificial Intelligence*, volume 36, pages 4637–4645, 2022.

77. T. Yatsunenko, F. E. Rey, M. J. Manary, I. Trehan, M. G. Dominguez-Bello, M. Contreras, M. Magris, G. Hidalgo, R. N. Baldassano, A. P. Anokhin, et al. Human gut microbiome viewed across age and geography. Nature, 486(7402):222–227, 2012.

78. Z. Ying, J. You, C. Morris, X. Ren, W. Hamilton, and J. Leskovec. Hierarchical graph representation learning with differentiable pooling. Advances in neural information processing systems, 31, 2018.

79. Z. Zhang, J. Geng, X. Tang, H. Fan, J. Xu, X. Wen, Z. Ma, and P. Shi. Spatial heterogeneity and co-occurrence patterns of human mucosal-associated intestinal microbiota. The ISME journal, 8(4):881–893, 2014.

80. Z. Zhang, P. Zhao, X. Li, J. Liu, X. Zhang, J. Huang, and X. Zhu. Signed graph representation learning: A survey. *arXiv preprint arXiv:2402.15980*, 2024.

81. Y. Zhou and F. Zhi. Lower level of bacteroides in the gut microbiota is associated with inflammatory bowel disease: a meta-analysis. BioMed research international, 2016(1):5828959, 2016.

